# A global timing mechanism regulates cell-type specific wiring programs

**DOI:** 10.1101/2020.09.18.304410

**Authors:** Saumya Jain, Ying Lin, Yerbol Z. Kurmangaliyev, Javier Valdes-Aleman, Samuel A. LoCascio, Parmis Mirshahidi, Brianna Parrington, S. Lawrence Zipursky

## Abstract

The assembly of neural circuits is dependent upon precise spatiotemporal expression of cell recognition molecules^1–6^. Factors controlling cell-type specificity have been identified^7–9^, but how timing is determined remains unknown. Here we describe the induction of a cascade of transcription factors by a steroid hormone (Ecdysone) in all fly visual system neurons spanning target recognition and synaptogenesis. We demonstrate through single cell sequencing that the Ecdysone pathway regulates the expression of a common set of targets required for synaptic maturation and cell-type specific targets enriched for cell surface proteins regulating wiring specificity. Transcription factors in the cascade regulate the expression of the same wiring genes in complex ways, including activation in one cell-type and repression in another. We show that disruption of the Ecdysone-pathway generates specific defects in dendritic and axonal processes and synaptic connectivity, with the order of transcription factor expression correlating with sequential steps in wiring. We also identify shared targets of a cell-type specific transcription factor and the Ecdysone pathway which regulate specificity. We propose neurons integrate a global temporal transcriptional module with cell-type specific transcription factors to generate different cell-type specific patterns of cell recognition molecules regulating wiring.

---

Animal behavior is dependent upon the formation of neural circuits with high fidelity. Many cell-surface proteins, particularly of the Immunoglobulin (Ig), Leucine-Rich Repeat and Cadherin families, promote interactions between neurons during circuit assembly^1, 2^. How does a limited set of cell-surface protein coding genes (∼10^3^ genes in fruit fly^10^) specify a much larger number of synapses (∼10^7^ synapses amongst ∼10^5^ neurons in fly brain^11^)? This is made possible by the use of several developmental strategies including the reuse of the same molecules to determine multiple, spatially and temporally separated neuron-neuron interactions^3, 4, 6, 12^. Wiring specificity, thus, relies on genetic programs which regulate the cell-type and temporal specificity of genes encoding recognition molecules. Many programs that define cell-type specificity have been described^7–9^. Here we set out to discover how temporal specificity is determined.

## Dynamic expression of wiring genes

We defined metrics for cell-type specificity and temporal dynamics for each gene using our previously reported transcriptomic dataset of 97 neuron-types in the fly visual system across seven developmental time points (24 – 96 hours after pupal formation (hAPF))^12^ (Fig. 1a). This period encompasses dendritic and axon terminal morphogenesis within target regions, selection of appropriate synaptic partners and synaptogenesis. We assigned two scores to each gene: 1) Cell-type variability score, which provides a measure of cell-type specificity; and 2) Temporal dynamicity score, which provides a measure of changes in expression across developmental time (see Methods, Fig. 1a, Extended data Table 2).

**Fig. 1.**
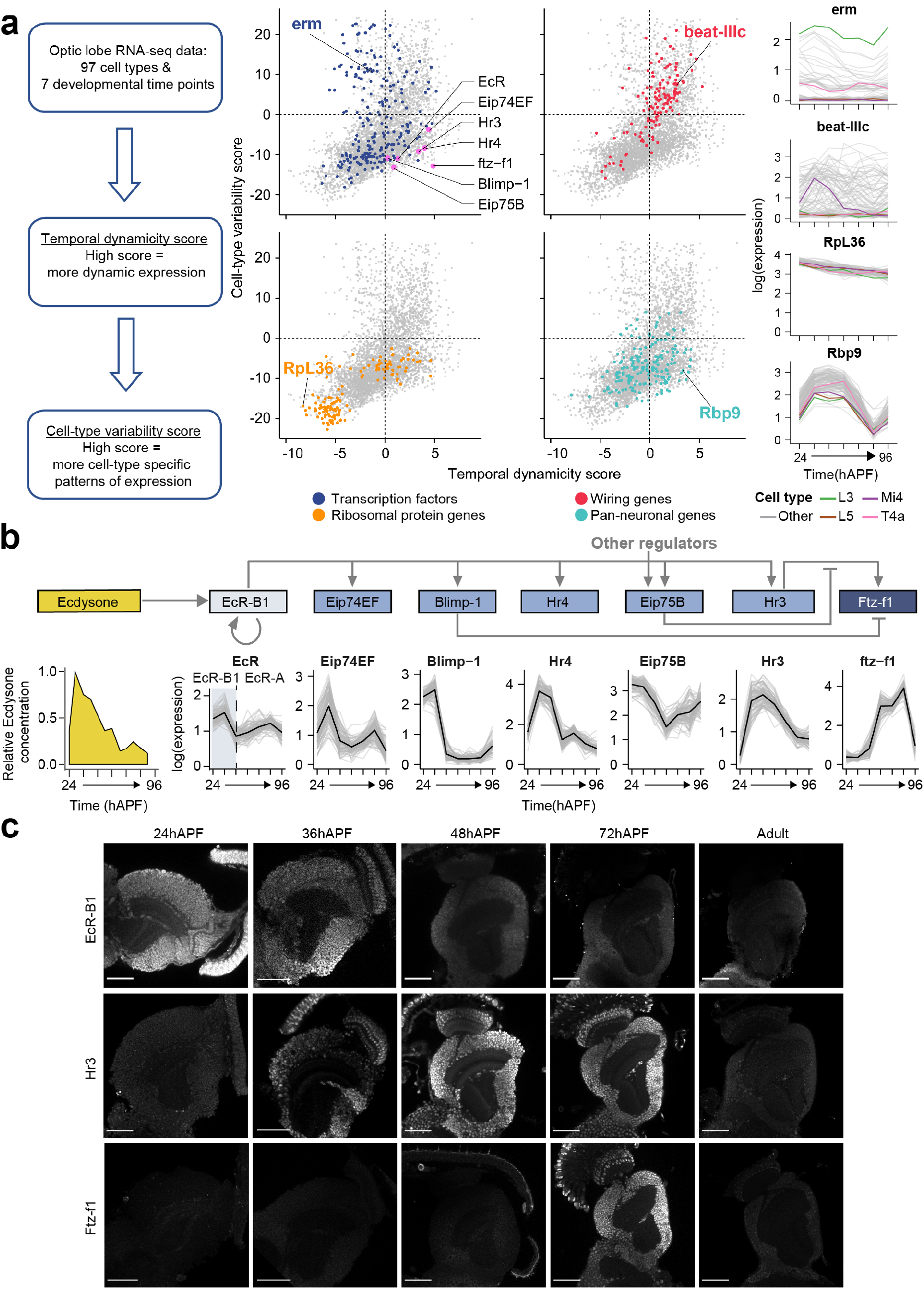
Dynamic expression of wiring genes and Ecdysone pathway TFs. **a,** Plots showing Cell-type variability and Temporal dynamicity scores calculated using data from Kurmangaliyev *et al.*, 2020^12^. Each plot highlights genes belonging to the indicated gene group. Right, expression of one gene from each group, highlighting expression in L3, L5, Mi4, T4a neuron-types. Transcription factors of the Ecdysone-pathway are highlighted (pink) in top-left plot. **b,** Ecdysone-pathway showing a subset of known genetic interactions. Relative concentration of Ecdysone over development is shown (adapted from Pak and Gilbert, 1987^19^). Line plots: each grey line is expression in one neuron-type. Black line is mean across all neuron-types. Isoform change from EcR-B1 to EcR-A is highlighted (also see Extended data Fig. 1). All expression plots: time in hours after pupa formation (hAPF). All expression is log expression values from Kurmangaliyev *et al.* **c,** Images showing immunostaining for EcR-B1, Hr3 and Ftz-f1 at the indicated time points. Scale bar, 50μm.

“Housekeeping” genes, such as ribosomal protein coding genes (e.g. RpL36), generally have low scores on both scales, suggesting low variation in expression across cell-types and time. Many transcription factors (TFs) display high cell-type variability but low dynamicity scores (e.g. erm), consistent with their role in establishing and maintaining neuron types^6, 12, 13^. A previously identified set of genes representing a pan-neuronal program demonstrate low variability scores and a range of dynamicity scores (e.g. RNA-binding protein, Rbp9)^12^. Finally, we compiled a list of cell surface molecules (CSMs) with previously described roles in wiring (and paralogs thereof), into a group we hereafter refer to as ‘wiring genes’ (see Methods, Extended data Table 3), and found that they have high cell-type variability and temporal dynamicity scores (e.g. beat-IIIc). Dynamic, cell-type specific expression is consistent with previous reports for expression patterns for these genes^5, 6, 12^. While cell-type specific TFs have been identified which control wiring genes, what controls their temporal specificity remains unknown.

## A cascade of nuclear receptors is expressed in visual system neurons

To identify candidates for temporal regulators, we focused on TFs with high dynamicity scores (Fig. 1a, Extended data Table 2). These were enriched for nuclear receptors (GO term: “nuclear receptor activity”, p-value 9.3 × 10^-5^; Reactome pathway analysis: “Nuclear Receptor transcription pathway” p-value 8.3 × 10^-10^; also see Extended data Table 4). Nuclear receptor TFs (such as Hr3, Hr4 and ftz-f1) which exhibit high temporal dynamicity (Fig. 1a), are components of a cascade of TFs activated by the steroid hormone Ecdysone (Fig. 1b). Indeed, the Ecdysone Receptor (EcR) itself, as well as other TFs in this cascade are expressed in a synchronous way in all neurons in the developing visual system (at the mRNA and protein level, Fig. 1b, c, Extended data Fig. 1). This includes a synchronous change in isoform from EcR-B1 to EcR-A^14^. The timing and order of expression of these TFs correlates with a mid-pupal pulse of Ecdysone^15–19^ (Fig. 1b). Consistent with control of dynamic gene expression, ATAC-Seq^20^ at 40hAPF, 60hAPF and 72hAPF in L1 neurons (see Methods) identified an enrichment for predicted binding sites for EcR, Hr3 and Ftz-f1 in regions with dynamic patterns of chromatin accessibility^21^ (Extended data Fig. 2, Extended data Table 5). Although these TFs are expressed in all neurons (Fig. 1b, c), they are known in other contexts to regulate a cell-type specific set of genes^22^. These observations raised the possibility that the Ecdysone-pathway has been co-opted as a temporal regulator to control the dynamic and cell-type specific expression of wiring genes.

## EcR controls neurite terminal morphology and synapse distribution

To test the role of the Ecdysone-pathway in wiring, we sought tools available for disrupting the pathway. Given the proximity of the EcR gene to the centromere and the lethality of whole animal mutants, phenotypic analysis using null-mutants is difficult to perform across cell-types (see mosaic analysis with *EcR* mutants in L5 below). We thus, began by expressing a widely used dominant-negative allele of EcR, EcR^DN^ (mutant with ligand-binding defect^23^, EcR-B1^W650A^), in a range of neuron-types arborizing in different neuropils and layers using the GAL4-UAS system (Fig. 2a, Extended data Fig. 3). Targeted expression of EcR^DN^ led to profound defects in axonal and/ or dendritic branching across all neuron-types tested, namely lamina neurons (L1-L5), T4/T5, Mi4 and Dm4 (Fig. 2, Extended data Fig. 3). EcR^DN^-induced defects in L1-L5 arborization were also seen with EcR-RNAi and could be largely rescued by overexpression of EcR-B1 from a cDNA transgene (Extended data Fig. 3a).

**Fig. 2.**
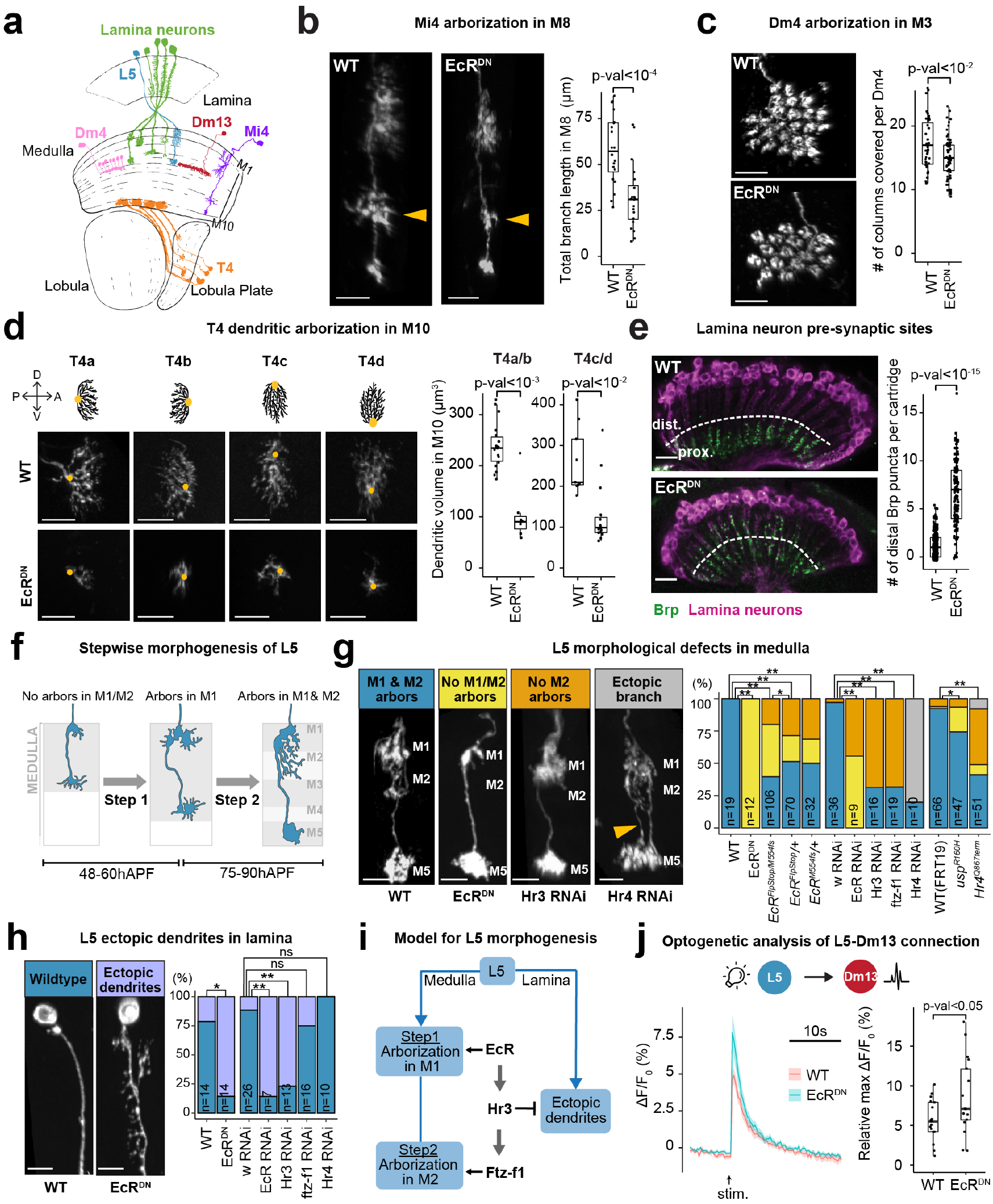
Ecdysone-pathway TFs control multiple aspects of wiring. **a,** Schematic of Lamina (L1-L5), Mi4, Dm4, Dm13 and T4. **b,** Morphology of Mi4 ± EcR^DN^. Yellow arrowhead: M8 layer. Scale bar, 10µm. Total branch length in M8 is quantified. (WT, 20 neurons, 3 animals; EcR^DN^, 20 neurons, 4 animals). **c,** Morphology of Dm4 ± EcR^DN^. Total number of columns covered/ neuron is quantified. (WT, 39 neurons, 5 animals; EcR^DN^, 61 neurons, 5 animals). **d,** Schematic of dendrites of T4 a-d subtypes. Dorsoventral (D, V) and anterior-posterior (A, P) axes are shown. Below, dendrite morphology ± EcR^DN^. Orange dot, dendritic stalk. Scale bar, 10µm. Dendritic volume for T4a, b and T4c, d subtypes is quantified. (WT, 18 T4a/b, 9 T4c/d neurons, 4 animals; EcR^DN^, 7 T4a/b, 15 T4c/d neurons, 5 animals). **e,** Lamina neuron (magenta) presynaptic sites (Brp, green) in the lamina neuropil ± EcR^DN^. Scale bar, 50µm. Distal Brp puncta/ cartridge is quantified. (WT, 409 cartridges, 7 animals; EcR^DN^, 153 cartridges, 6 animals.) **f,** Schematic of stepwise morphogenesis of L5 arbors. **g,** L5 arborization defects and their distributions under the given genotypes (also see Extended data Fig. 5a). **h,** L5 ectopic dendrite phenotype with quantification under given genotypes. **g, i,** n = number of neurons (all conditions, animals ≥ 4). **g, h,** Scale bar, 5µm. **i,** Schematic showing roles for EcR, Hr3 and Ftz-f1 in L5 branching. **j,** GCaMP6s response in Dm13 upon optogenetic stimulation of WT and EcR^DN^-expressing L5 neurons. Amplitude of peak response for each condition is given. (WT, 17 animals; EcR^DN^, 17 animals.) **b -e, j,** p-values (two-sided Student’s t-test) are given. **g, h,** *, p-value (one-sided Fisher’s exact test) < 0.05, ** p-value < 0.001.

We also observed that expression of EcR^DN^ in lamina neurons led to a redistribution of their presynaptic sites in the lamina neuropil from strictly proximal to more distal regions of the lamina (Fig. 2e). *DIP-β* mutants have been shown to produce a similar phenotype^24^, and via scRNA-Seq experiments described below, we identified DIP-β as a target of EcR in L4 neurons (Extended data Fig. 4a, b). That DIP-β is directly regulated by EcR, is consistent with the presence of a strong EcR binding site within the first intron of DIP-β (see Methods, Extended data Fig. 4c). Thus, EcR controls dendritic and axonal terminal morphology and presynaptic site distribution across cell-types in the lamina, medulla and lobula plate.

## Different EcR pathway TFs regulate specific steps in L5 wiring

We next sought to investigate roles for the EcR-pathway at different steps in wiring of a single neuron-type. L5 axon terminals target to medulla layer M5 during early pupal development^25, 26^, then arborize in M1 (between 48hAPF and 60hAPF), followed by extension of these arbors from M1 to M2 (between 75hAPF and 90hAPF) (Fig. 2f).

Targeted expression of EcR^DN^ in developing L5 neurons did not disrupt targeting to the M5 layer, which is completed prior to the mid-pupal Ecdysone pulse. However, it led to a complete lack of branches in M1/M2 (Fig. 2g, Extended data Fig. 5a). To verify the role of EcR in establishing M1/M2 arbors, we generated a conditional null-allele for EcR using the FlpStop technology^27^ (EcR^FlpStop^, see Methods, Extended data Fig. 5b). Similar to EcR^DN^, we saw loss of M1/M2 arbors in: 1) EcR^FlpStop^ over a frameshift mutation in the ligand binding domain (EcR^FlpStop/M554fs^); 2) Heterozygous L5 neurons (EcR^FlpStop/+^ or EcR^M554fs/+^); and 3) knock-down of EcR via RNAi. EcR mutants were phenocopied by a homozygous hypomorphic allele of *usp* (usp^R160H^). This gene encodes a cofactor which functions with EcR in different developmental contexts^28, 29^ (Fig. 2g). By contrast to its role in promoting branching in M1/M2 medulla layers, EcR suppresses L5 branching in the lamina (Fig. 2h). Wildtype L5 neurons (unlike L1-L4 neurons) do not elaborate branches in the lamina^30^. However, in response to either EcR RNAi or EcR^DN^, robust ectopic branching (similar in appearance to normal L1-L3 dendrites) was observed in the lamina.

While disruption of EcR activity blocks step 1 and step 2, knock-down of TFs downstream of EcR, Hr3 or Ftz-f1, selectively disrupts step 2 (Fig. 2g, Extended data Fig. 5a). These findings are consistent with EcR regulating the expression of genes that promote extension of arbors into M1 and that Hr3 and Ftz-f1 execute a distinct developmental step ∼ 24 hours later (Fig. 2g), promoting extension of arbors from M1 into M2. As Hr3 directly regulates *ftz-f1*^16^ (Fig. 1b, see Extended data Fig. 10), Ftz-f1 may act alone or in combination with Hr3, to regulate the expression of genes required for branching in M2. Loss of Hr4, another TF directly downstream of EcR, results in ectopic branches terminating in M5 (Fig. 2g, Extended data Fig. 5a). By contrast to the medulla, suppression of branching in the lamina relies on Hr3, but not Ftz-f1 (Fig. 2h, i).

To see if the Ecdysone-pathway also affects communication between L5 and its postsynaptic partners, we used an L5-specific Gal4 to express Chrimson with or without EcR^DN^ and recorded Ca^2+^ responses in different postsynaptic cells (using GCaMP6s) upon activation of Chrimson by light. Given the near complete loss of M1 arbors with EcR^DN^, we focused on L5 postsynaptic partners in the M5 layer, Dm13 and Tm3^31^. Optogenetic stimulation of EcR^DN^-expressing L5 neurons led to a stronger response in Dm13 (Fig. 2j), while responses from Tm3 remained unchanged (Extended data Fig. 6a). A postsynaptic neuron-type specific change in GCaMP6s response is consistent with perturbation of L5 synaptic connectivity upon disruption of the Ecdysone-pathway.

In summary, the Ecdysone-pathway sculpts L5 connectivity in multiple ways during development, including affecting branches at different times, in different ways and within distinct subcellular domains (see summary in Figure 2i).

## The EcR cascade regulates pan-neuronal and cell-type specific genes

To identify genes regulated by the EcR cascade, we focused on lamina neurons (L1-L5). These were enriched by Fluorescence-activated cell sorting^32^ using a pan-lamina marker and profiled using a single-cell RNA-Seq (scRNA-Seq) based approach similar to the one used in Kurmangaliyev *et al*^12^. Transcriptional profiles of wildtype lamina neurons determined this way were similar to the ones determined from dissociated whole optic lobes^12^ (Extended data Fig. 7). We then profiled the transcriptomes of EcR^DN^, EcR RNAi or Hr3 RNAi-expressing lamina neurons and controls by scRNA-Seq at four timepoints (24hAPF, 48hAPF, 72hAPF and Adult, see Extended data Fig. 7a)^12^.

By comparing the transcriptomes of wildtype and EcR^DN^-expressing lamina neurons, we identified 872 downregulated and 517 upregulated genes in EcR^DN^-expressing cells (fold change > 2, p-value < 0.05) (Fig. 3a; Extended data Fig. 8). Genes affected by EcR^DN^ are similar to the set of genes affected by EcR RNAi, though the latter is significantly less effective at blocking the Ecdysone pathway in this context (Extended data Fig. 9, 11a, c, e). Hr3 RNAi also has a similar effect on gene expression as EcR^DN^, especially at later time points (Extended data Fig. 10, 11b, d, e) consistent with the genetic relationship between EcR and Hr3 (Fig. 1b). In contrast to L1 and L3-L5, EcR^DN^ had modest effects on gene expression in L2, consistent with lower EcR^DN^ expression in L2 neurons at 48hAPF using the pan-lamina GAL4 driver (Extended data Fig. 8b – d, f). Thus, the EcR pathway regulates hundreds of genes in developing lamina neurons.

**Fig. 3.**
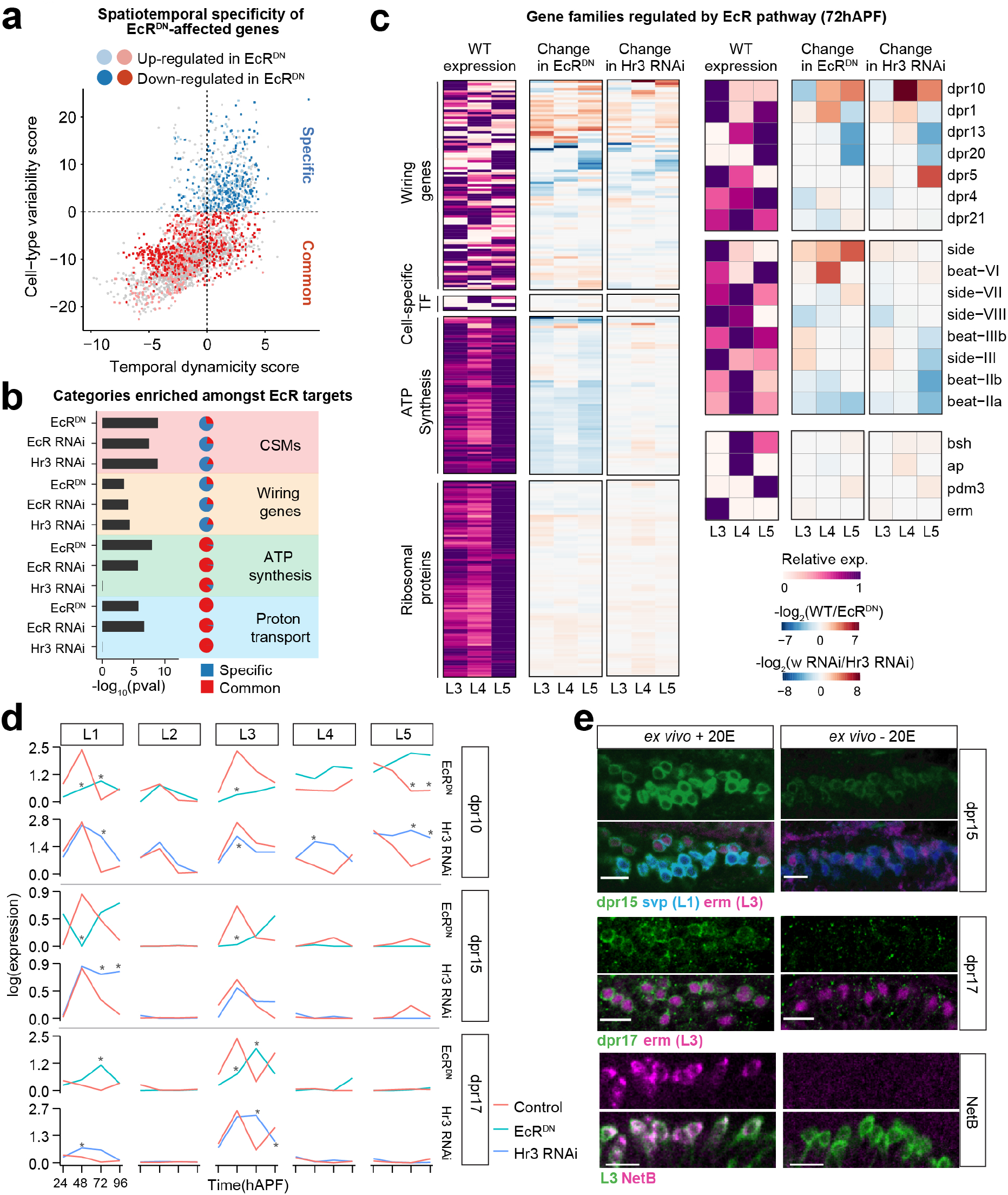
Common and cell-type targets of the Ecdysone-pathway. **a,** Cell-type variability vs Temporal dynamicity plot for EcR^DN^-affected genes (fold change > 2, p-value < 0.05). Cell-type Specific (blue) and Common (red) targets are shown. Darker colors, genes reduced in EcR^DN^; lighter colors, genes increased in EcR^DN^ (also see Extended data Fig. 11c, d). **b,** Gene groups enriched amongst genes reduced in EcR^DN^. Fraction of targets common to all neurons (red) and neuron-specific (blue) targets in each category are shown as ratio of red to blue in small pie charts. **c,** Relative WT expression at 72 hAPF (left), change in expression with EcR^DN^ (center), and change in expression with Hr3 RNAi (right) shown as heat maps for all genes expressed in L3-L5 neurons belonging to the specified gene categories at 72hAPF (also see Extended data Fig. 13a). Examples of wiring genes (e.g. Dpr family, Side-Beat family) and cell-type specific transcription factors (e.g. erm) are shown separately. **d,** dpr10, dpr15, dpr17 expression ± EcR^DN^ (above) and ± Hr3 RNAi (below). * p-value < 0.05, fold change > 2. Expression is in log values. **e,** Staining for dpr15 and dpr17 reporters (MiMIC lines, see Methods), and NetB (using anti-NetB antibody) in lamina neuron cell-bodies ± 20 Hydroxyecdysone (20E – active form of Ecdysone) in the medium (also see Extended data Fig. 14) in *ex vivo* preparations of pupal brains (see Methods). Note most NetB protein accumulates in L3 growth cones in the M3 layer (not shown).

Analyses of the targets of the Ecdysone-pathway revealed that this pathway: 1) can both activate and repress gene expression (Fig. 3a, c, d, Extended data Fig.12); 2) does not affect cell-fate (Extended data Fig. 8e, 9e, 10e); and, 3) controls a larger number of genes with cell-type specific, dynamic expression than expected by chance (one-sided Fisher’s exact test, p-value = 0.005, Fig. 3b). Together, these observations establish that the Ecdysone-pathway controls different sets of genes expressed in specific cell-types in a dynamic fashion. Importantly, we found many dynamic genes to not be affected by EcR mutants, suggesting roles for other temporal regulators as well (see Discussion, Fig. 3a, Extended data Fig. 11c, d, 12d).

Gene ontology analysis revealed that targets of the EcR pathway were enriched for genes required for mitochondrial ATP biogenesis and neurotransmitter loading into synaptic vesicles (see ‘proton transport’, Fig. 3b, c, Extended data Fig. 13b, Extended data Table 2), processes required for synapse development, function or both^33–36^. Importantly, they have low cell-type variability scores, and are thus likely regulated by the Ecdysone-pathway across neuron-types (Fig. 3b, also see common targets, Fig. 3a).

## Control of cell-type specific wiring genes

The targets of EcR are also enriched for genes encoding CSMs, including wiring genes (Fig. 3b, c). As opposed to genes required for ATP-synthesis and vacuolar ATPases, CSMs and wiring genes regulated by the Ecdysone-pathway are largely composed of targets with cell-type specific expression (Specific targets, Fig. 3a, b). These genes can be repressed in one cell-type and activated in another (see dpr17 in L1 and L3, Fig. 3d), and can be regulated at different times in different cell types (e.g. dpr10 in L1 and L5). In some cases, bidirectional changes in gene expression resulted from activation by EcR, followed by repression by Hr3, indicative of a form of feedback inhibition (e.g. see dpr10 and dpr15 in L1 and dpr17 in L3, Fig. 3d). Thus, activation and repression by different Ecdysone-pathway TFs control cell-type and temporally specific patterns of gene expression of wiring genes.

To assess if genes encoding CSMs affected by disruption of EcR activity are responsive to hormone, we looked at the expression of some of these genes by controlling availability of Ecdysone in *ex vivo* preparations. We modified a protocol developed by Özel *et al*, wherein brains are removed from pupae (at 22hAPF) prior to the mid-pupal Ecdysone pulse, and then incubated in media with or without the active form of Ecdysone, 20 Hydroxyecdysone^37^ (20E, Extended data Fig. 14a, see Methods). Consistent with scRNA-Seq data, the expression of several targets of EcR depends upon inclusion of ligand in the media (Fig. 3e, Extended data Fig. 14b).

Together, these findings indicate that the Ecdysone-pathway, expressed in a common temporal fashion in all neurons, regulates genes required for proper circuit formation in different cell types in temporally specific ways.

## Erm and EcR co-regulate wiring genes in L3

We next sought evidence that cell-type specific wiring programs are co-regulated by cell-type specific and EcR pathway TFs. Here, we focused on a conserved TF, Erm (Fezf1/2 in mammals) which is expressed in post-mitotic L3 neurons throughout development, but not other lamina neurons, and controls several L3-specific wiring programs^32, 38^. Though Erm is expressed in a largely stable fashion throughout development, many of its known targets^38, 39^ (including wiring genes) show dynamic expression (Fig. 4a). Interestingly, genes co-regulated by Erm and EcR are enriched for dynamic genes (one-sided Fisher’s exact t-test, p-value = 0.0331, see Methods). Additionally, EcR pathway TFs are expressed sequentially in developing L3 neurons. Together, these observations are consistent with Erm and the Ecdysone-pathway acting together to determine the spatiotemporal expression patterns of wiring genes in L3.

**Fig. 4.**
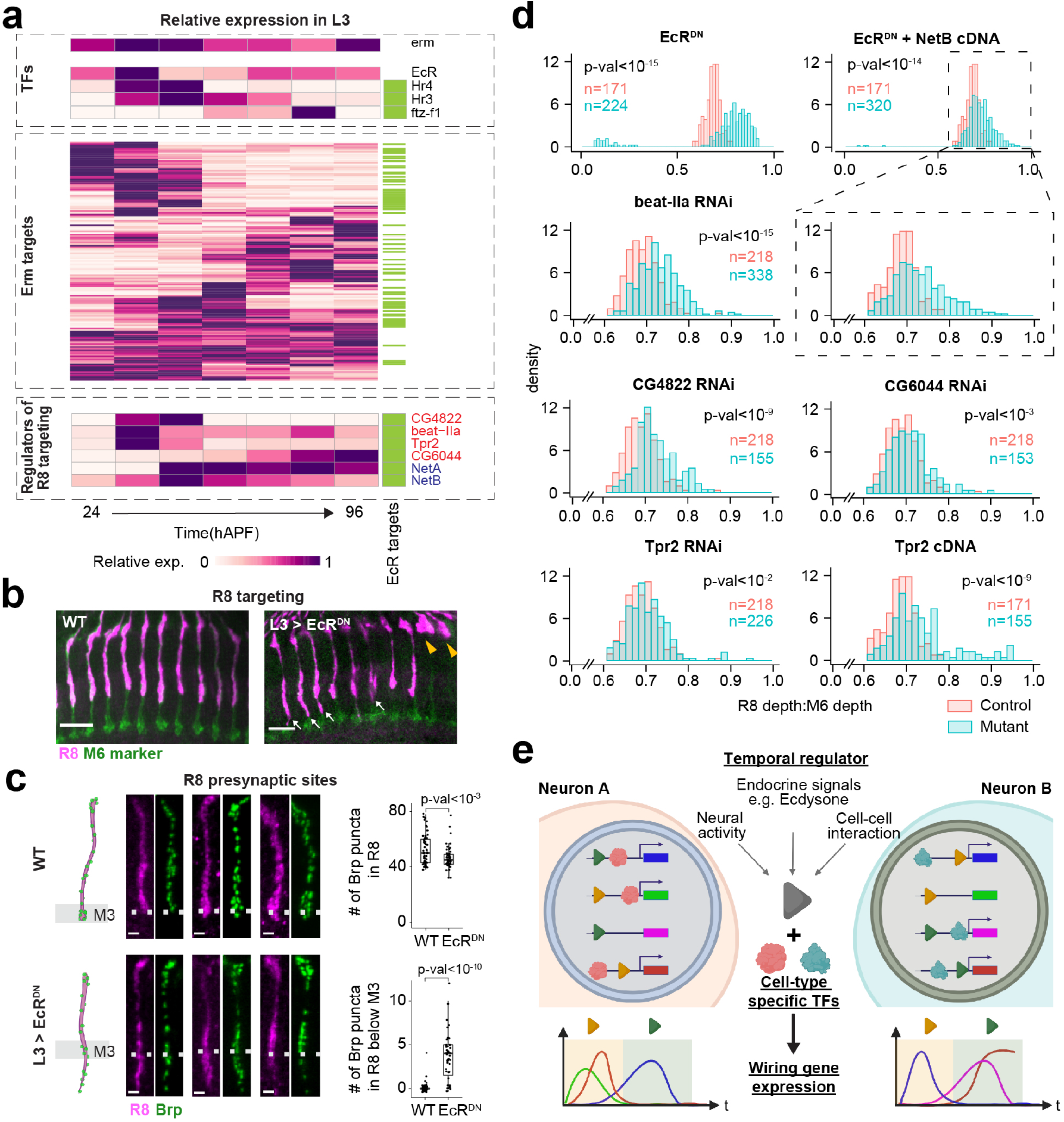
Screening EcR and Erm co-regulated genes for wiring regulators. **a,** Heatmap showing relative expression of erm, EcR-pathway TFs and Erm targets in L3 neurons^38, 39^. Green box indicates genes affected by EcR^DN^ in L3 neurons. Genes that regulate R8 targeting are shown below as indicated. NetA and NetB (blue) are known regulators. Genes in red are identified in RNAi screen in this study. **b,** R8 axons ± EcR^DN^ expression in L3. Magenta, R8 neurons; green, M6 marker (m24B10); yellow arrowheads, R8 axons terminating short of M3 and; white arrows, R8 terminals extending beyond M3. Scale bar, 10µm. **c,** R8 presynaptic sites (Brp, green) ± EcR^DN^ expression in L3. Right, quantification of Brp puncta number under the conditions shown. Scale bar, 5µm. (WT, 58 neurons, 3 animals; EcR^DN^, 47 neurons, 3 animals). **d,** Distributions of R8 axon terminal depth (as in **b**) in control (red) or with expression of RNAi or cDNA (as shown, blue) in L3 neurons. Numbers of neurons/ condition are given. All conditions, number of animals ≥ 3. p-values (two-sided Student’s t-test in **c,** Kolmogorov-Smirnov test in **d**) are given. **e,** Model for cell-type specific temporal control of wiring gene expression (see text).

L3 neurons non-cell autonomously control the targeting of R8 neurons to the M3 layer. Netrin, which requires both Erm^38^ and EcR for proper dynamic expression in L3 (Fig. 3e, Extended data Fig. 16), is required for adhesion to the M3 layer, but not for recognition of the layer as its target. Indeed, in *netrin* mutants R8s target to the appropriate later but then retract to distal layers of the medulla^40^. While some R8s retract to distal medulla in *erm* and *EcR* mutants, most terminate beyond M3^38^ (Fig. 4b, d, Extended data Fig. 15b). This is consistent with Erm and EcR acting together to control M3 target layer recognition and subsequent netrin-dependent stabilization. R8 terminals extending into deeper layers also contain ectopic presynaptic sites and show a modest overall decrease in synapse number (Fig. 4c, Extended data Fig. 15a), consistent with defects in R8 synaptic connectivity. We thus speculated that in addition to netrin, other genes co-regulated by EcR and Erm (see Extended data Fig. 16) control R8 targeting.

To identify regulators of R8 targeting, we performed an RNAi screen for 12 genes co-regulated by EcR and Erm (Fig. 4a, d, Extended data Fig. 15b, c, Extended data table 7, 8). Of these, RNAi for beat-IIa, CG4822, CG6044 and Tpr2, led to R8 targeting past the M3 layer (Fig. 4d). Beat-IIa and CG6044 are members of the Ig superfamily, a class of genes known to mediate cell recognition^1, 2^. Tpr2 regulates the activity of steroid hormone receptors in mammals, and may play an analogous role in regulating EcR-pathway TFs. Indeed, as in mammals where either overexpression or knock-down of Tpr2 results in reduced hormone receptor activity^41, 42^, Tpr2 overexpression in L3 neurons phenocopies Tpr2 RNAi (Fig. 4d). And finally, CG4822, which belongs to a subfamily of ABC transporters involved in cholesterol and steroid efflux, is also implicated in steroid regulation^43, 44^.

Together these data indicate that the EcR-pathway and Erm co-regulate the spatiotemporal specificity of several genes in L3 required for proper R8 wiring, including genes regulating steroid abundance or steroid receptor function (CG4822 and Tpr2), genes encoding Ig superfamily proteins (Beat IIa and CG6044) and secreted proteins (NetA and B) required for targeting to and stabilization of R8 growth cones in M3.

## Discussion

Wiring an animal brain requires precise cell-type and temporally specific expression of cell recognition proteins^3–5, 32, 45^. The Ecdysone-pathway functions as a temporal regulator of major developmental transitions, including larval molts and metamorphosis, as well as controlling sequential divisions of postembryonic neuroblasts and neuronal re-modelling^15, 22, 46–48^. Here we demonstrate that the EcR cascade has been co-opted for the temporal regulation of neuron-type specific wiring programs within the developing neuropil. Neuron-type specificity in this case is, at least in part, determined by the activity of cell-type specific TFs. The ability of the EcR cascade to function as a flexible temporal regulatory module may be similar to modules regulating spatial patterning (e.g. Hox genes and BMP/Wnt/Hh pathways^49–51^) which act in context-dependent ways to generate different morphological outcomes in many different developmental contexts.

Global pathways, including other steroid and peptide hormones, waves of stimulus independent neural activity, sensory experience as well as local intercellular interactions contribute to wiring more broadly in controlling temporal aspects of development in the vertebrate and invertebrate CNS (Fig. 4e)^52–57^. As in the EcR pathway in the fly visual system, these temporal signals may also alter the activity of broadly expressed TFs (e.g. via phosphorylation) which combine with cell-type specific TFs to control the expression of cell surface molecules regulating synaptic connectivity.

## METHODS

### Fly husbandry and stocks

Maintenance and rearing of fly lines, as well as staging of pupae was done as described in Tan *et al.*^32^ Fly lines used in this work are listed in Supplementary Table 1.

### Data analyses of fly optic lobe RNA-seq

#### Measure of cell-type variability and temporal dynamicity for each gene

RNA-seq data of developing fly optic lobe were generated by Kurmangaliyev *et al.*^12^, which contains 97 cell types covering 7 time points. For each gene, we assigned a cell-type variability score by their cell-type specific expression and a temporal dynamicity score by their expression changes over development. Cell-type variability score was calculated by the variance of gene expression at each time point when the gene is expressed, and then the maximum variance was normalized by the average expression across cell types that time point. Temporal dynamicity score was calculated by the sum of the absolute changes between consecutive time points for each cell type where the gene is expressed, and then the averaged value across cell types is normalized by the average of their peak expression. Both scores were centered their mean to 0 and transformed to natural log for visualization.

#### Wiring genes

Genes were designated to the category of ‘wiring genes’ if they satisfied the following criteria: 1) they were amongst the top 50% of the highest expressed cell-surface or secreted molecules in the developing optic lobe (from Kurmangaliyev *et al.*), 2) either themselves, or their paralogs (DIOPT v8.0 score ≥ 4) had published roles in establishing neuronal morphology or connectivity in *Drosophila*. Extended data Table 3 contains a list of wiring genes along with references to published studies describing their roles in wiring.

#### Gene Ontology and REACTOME analysis of dynamic TFs

Dynamic TFs were defined as ones with temporal dynamicity score > 0 (see Fig. 1a, Extended data Table 2). Gene ontology analysis was performed using GOrilla (http://cbl-gorilla.cs.technion.ac.il) using dynamic TFs ranked on the basis of temporal dynamicity scores. All enriched categories are given in Extended data Table 4. For REACTOME pathway analysis, dynamic TFs were entered into the ‘Analyze Data’ tool on reactome.org (https://reactome.org/PathwayBrowser/#TOOL=AT). Reactome pathway analysis for *Drosophila melanogaster* was then run with default parameters.

### Bulk RNA-seq

Bulk RNA-Seq analysis of L1 neurons at 40hAPF, 60hAPF and 72hAPF was performed as in Tan *et al.*^32^ At least 2 replicates were generated for each timepoint. A minimum of 2000 cells were sorted for each experiment. cDNA libraries were created using the SMART-Seq2 protocol^58^, which were then sequenced together using the HiSeq4000 platform (paired-end 50bp). Raw fastq reads files were mapped to FlyBase reference genome (release 6.29) using STaR (2.6.0) and only uniquely mapped reads were collected. Genes with counts per million (CPM) ≥ 4 in more than 2 samples were used for normalization with R package edgeR (3.26.8). Gene expression was quantified using RPKM units (Reads Per Kilobase of exon per Million reads mapped), calculated based on reads in the sum of exons using customized scripts. The correlation between biological replicates were calculated using Spearman correlation for top 500 genes with the highest variation across all samples. Coverage tracks were generated by bamCoverage(3.5.1).

### Bulk ATAC-seq

At least 8000 L1 neurons were FACS purified at 40hAPF, 60hAPF and 72hAPF as described in Tan *et al.*^32^ Biological duplicates were generated for each timepoint. ATAC-Seq libraries were generated as per the following protocol, which was modified from the one described in Buenrostro *et al.*^59^: 1) FACS purified cells were collected in 300ul 1X PBS and spun at 800xg for 8’ at 4°C. 2) Cell pellet was then directly resuspended in 50µl of a tagmentation enzyme mix (25µl Nextera DNA library prep kit – Tagment DNA Buffer, 19µl Nuclease-free water, 5µl 1% IGEPAL CA630, 1µl Nextera DNA library prep kit - Tagment DNA Enzyme) and incubated at 37°C for 30’ with constant agitation at 400rpm. 3) DNA was purified from the tagmentation mixes using the Qiagen minElute Reaction Cleanup Kit as per the manufacturer’s protocol. 4) A 10µl qPCR reaction was setup to determine the optimum number of amplification 455 cycles (similar to Buenrostro *et al.*). 5) A larger PCR reaction (40µl) was then setup and PCR products in the 200-500bp range were gel purified. 6) Steps 4 and 5 were repeated, this time with barcoded primers (see Buenrostro *et al.*) to allow multiplexing of all libraries onto a single lane of HiSeq4000 (paired-end 50bp).

Raw fastq reads files were mapped to FlyBase reference genome (release 6.29) using Bowtie2 (2.2.9) and uniquely mapped genes were kept. All samples were pooled together prior to peak calling. Read start positions were shifted +4 or -5bp and used for peak calling using MACS2 (2.1.1) with parameters “-q 0.01 --nomodel --shift -100 --extsize 200”. There were 26,122 peaks identified. Then bedtools multicov (2.27.1) was used to sum the total reads within each peak separately for each sample. Peaks with CPM ≥ 4 were used for counts normalization with edgeR. Correlations between biological replicates were calculated using Spearman correlation for the top 500 peak regions with the highest variation of peak levels across all samples. Distribution of the top 5000 peaks across different genomic features at 40hAPF, 60hAPF and 72hAPF was determined using the R package ChIPseeker (1.20.0). Each ATAC peak was associated to the nearest gene based on proximity to the transcription start site using ChIPseeker. To compare change of ATAC-seq peak accessibility over time and change in expression of their nearest gene, log_2_(fold change ATAC-seq peak read coverage) was plotted against log_2_(fold change expression of nearest gene) for dynamic ATAC-Seq peak (Extended data Fig. 2b). Bulk RNA-Seq data was used for this, as this transcriptome dataset had been generated for the same time points as for ATAC-Seq. This was done separately for peaks differentially accessible between 40hAPF and 60hAPF (Extended data Fig. 2b (left)) and between 60hAPF and 72hAPF (Extended data Fig. 2b (right)). Differential peak analysis was performed using edgeR. Fold-change> 2 and adjusted p-value < 0.05 were chosen as the cutoffs to define peaks with differential accessibility between time points (dynamic peaks).

Binding sites for the EcR-usp complex (motif: Jaspar MA0534.1), Hr3 (motif: Fly Factor Survey, Hr46_SANGER_5) and Nr5a1, mammalian homolog of Ftz-f1 (motif: Jaspar 1540.1) across the genome were identified using the PWMscan (https://ccg.epfl.ch/pwmscan/). Cutoff of p-value < 10^-5^ were used to identify binding sites^60^. Enrichment of binding sites within different sets of ATAC-Seq peaks (Extended data Fig. 2d) was done using Homer (v4.7) mergePeaks utility. p-value ≤ 0.01 was considered as significantly more overlap than expected by chance.

### Calcium imaging

Calcium imaging experiments were performed using a 3i VIVO Multiphoton upright microscope (Intelligent Imaging Innovations) with a water immersion objective (W Plan-Apochromat 20x/1.0 DIC, Zeiss). L5 neurons expressing CsChrimson (using 64B07 Gal4) were photo-stimulated with a 1040 nm laser (SpOne-8, Spectra-Physics) coupled to a 2-photon Phasor (Intelligent Imaging Innovations) to generate a holographic pattern to restrict the stimulation area. GCaMP6s responses were recorded using an imaging laser tuned to 940 nm (Chameleon Discovery, Coherent).

Experimental crosses were set in vials with fly food supplemented with all-trans retinal (0.5 mM) and kept in the dark at 25 °C. The brains of ≤ 5-day old adult flies were dissected in a saline solution (135 mM NaCl, 5 mM KCl, 2 mM CaCl_2_.2H_2_O, 4 mM MgCl_2_.6H_2_O, 5 mM TES, 36 mM Sucrose, pH 7.15) and mounted dorsal side on a poly-L-lysine-coated coverslip. Typically, the dissected brains were held for about 5 minutes before the start of the experiment to ensure a stable basal calcium signal. The photo-stimulation hologram was restricted to a rectangular area of 18 μm x 65 μm to roughly cover the middle third of the axon layer of L5 neurons in the medulla layers 1-2. The experimental protocol consisted of three consecutive iterations of a 30 s resting period and a 500 ms stimulation event. The stimulation laser power at the objective end was 335 mW, measured with a power meter (PM100D Thorlabs). GCaMP responses were recorded from a single Z plane at 3.05 frames/s. This protocol was executed on both optic lobes when possible.

#### Image processing

The GCaMP image data were processed using custom macros in Fiji and analyzed using custom code written in R. Briefly, a region of interest (ROI) was manually defined as the medulla layers that include the sub-arbors of the neurons expressing GCaMP. The average pixel value inside such ROI was measured across all time points for each sample. All fluorescence values were reported relative to a fluorescence baseline (F_0_) defined as the median pixel value of the corresponding ROI during the entire imaging experiment. ΔF/F_0_ was calculated as ΔF/F_0_ = (Ft – F_0_)/F_0_, where Ft is the mean fluorescence value of the ROI at a given time point. The relative maximum ΔF/F0 was defined in a 10 s time window immediately after stimulation offset from which the recent baseline (mean ΔF/F_0_ of the 6.5 s preceding stimulation onset) was subtracted. Those failed individual trials in which there were no detectable responses were discarded. The three individual stimulation events from left and right hemispheres trials were averaged to generate a single response trace per animal.

### Multiplexed single cell transcriptomic analysis

For transcriptomic analysis of developing wildtype lamina neurons, w; UAS-H2A-GFP; 9B08-Gal4/Tm6B, tb females were crossed with males from different DGRP backgrounds (wildtype, see Supplementary Table 1 for list of DGRPs used). F1 prepupae were staged as in Kurmangaliyev *et al.*^12^ Males were only included in the analysis if no significant differences in gene expression were found with female pupae of the same genotype and developmental time point. Pupae corresponding to different developmental stages (see Extended data Fig. 7a) were all dissected, dissociated and processed at the same time. Each developmental time point was represented by ≥ 2 DGRP heterozygotes, and only one animal/ DGRP heterozygote was dissected. Tissue dissociation, FACS and preparation of single-cell libraries using 10X Genomics Chromium (v3) were carried out similar to Kurmangaliyev *et al.*, except that H2A-GFP expression was used to enrich for lamina neurons. All libraries were sequenced on a NextSeq500 platform (single-end 75bp).

For transcriptomic analysis involving UAS EcR^DN^ (BDSC #6872), males carrying a single DGRP autonomous chromosome – w; (UAS EcR^DN^ or UAS tdTom); DGRP/ Tm6b, tb were crossed with w; UAS-H2A-GFP; 9B08-Gal4/Tm6B females. For experiments involving UAS EcR RNAi (BDSC #9326) or UAS Hr3 RNAi (BDSC #27253), w; DGRP; (UAS wRNAi or UAS EcR RNAi or UAS Hr3 RNAi)/ Tm6B, tb males were crossed with UAS Dcr2; UAS H2A-GFP; 9B08 Gal4/ Tm6B, tb females (see Extended data Fig. 7a and Supplementary Table 1). UAS tdTom and UAS wRNAi were used as controls.

#### scRNA-seq data pre-processing

Raw fastq read files were processed using Cell Ranger (3.1.0) with default parameters. Seurat V3 was used for all preliminary analyses. FlyBase reference genome (release 6.29) was used for alignment and annotation. Profiled single cells were identified as coming from a particular developmental time-point and genetic background based on the natural genomic variance within the different parent DGRP lines. This was done using the pipeline described in Kurmangalyev *et al.*^12^, with the following modifications: the count of minor allele that equals to 2 among the analyzed DGRP strains were used as genomic variants to assign the profiled single cells to different DGRP parent lines. Only single cells with: 1) number of genes between 200 and 3000, 2) number of mitochondrial transcripts < 20% of all transcripts, and 3) assignment to a unique DGRP parent line; were used for all downstream analyses.

To identify different lamina neuron subtypes, all cells from a particular were integrated as described in Seurat v3 workflow and then subjected to unsupervised clustering, thus disregarding global temporal gene expression changes (similar to Kurmangaliyev *et al.*). Previously identified lamina neuron subtype specific genes^32^ were used to assign each cluster to a cell-type. All cell cluster separations are visualized in tSNE plots. Cells not assigned to a lamina neuron-type cluster were removed from subsequent analyses. Average expression for each gene for each cell-type at a particular timepoint and genetic background was calculated using normalized expression values prior to integration.

A global-scaling normalization used in Butler *et al.*^61^ is used in all scRNA-seq analysis in this paper. We normalize the gene expression measurement for each cell by the total expression and multiply by scale factor 10000. All expression shown in figures are natural log transformed.

### Comparison of scRNA-Seq with published dataset

For comparison of lamina neuron transcriptomes generated here with lamina neuron transcriptomes generated in Kurmangaliyev *et al.*, 500 genes with the highest variance across all cell types and all timepoints were used to calculate Spearman correlation.

### Differential gene expression analysis

Wilcoxon rank-sum test (fold change > 2, adjusted p-value < 0.05) was used to identify genes differentially expressed between timepoints, cell-types or as a consequence of EcR/ Hr3 perturbations. All fold change calculation used pseudo number 0.01.

### Enrichment analysis amongst EcR and Hr3 targets

For each of the four categories shown in Fig. 4b, enrichment was calculated using Fisher’s exact test. Briefly, number of genes expressed in lamina neurons overlapping with genes in the stated category vs other genes expressed in lamina neurons, was used to define a null hypothesis. Then genes affected by EcR^DN^, EcR RNAi or Hr3 RNAi were also divided into ones overlapping with genes in the stated category, and ones that don’t overlap. Number of genes in these two categories, along with similar numbers from the null-hypothesis, were then used to create a 2X2 matrix, which was then subjected to Fisher’s exact test.

Gene categories, Proton transport (proton transmembrane transport, GO:1902600) and ATP synthesis (ATP metabolic process, GO:0046034) were initially identified from GO analysis of targets of EcR^DN^ and EcR RNAi.

### Gene group analysis for EcR and Hr3 perturbation effect

Prior to clustering, all average gene expression from scRNA-seq transcriptomic analysis were normalized to maximum expression in each cell type across development. For each cell types, the relative expression of each gene was used to perform k-means clustering to assign expressed genes in different groups. The number of clusters for each cell type were determined by calculating Within-Clsuter of Squared Errors (WSS) and finding where the difference of WSS in consecutive cluster number first become less than 5. For Extended data Fig. 12, we presented the gene groups with the most up-regulated or the most down-regulated or the unchanged gene expression after expressing EcR^DN^ in each lamina neuron.

### *Ex vivo* culture of dissected brains

We used a protocol similar to the one described in Özel *et al.*^37^ Briefly, brains were dissected from 22hAPF pupae in pre-warmed Schneider’s medium (ThermoFisher #2172004) with either 1:1000 dilution of 1mg/ml 20-HydroxyEcdysone (dissolved in 100% Ethanol, Sigma H5142) or equivalent volume of 100% ethanol. Brains were then incubated in ∼ 200ul of medium ± 20-HydroxyEcdysone in 96 well plates for 26h at 25°C in a humidified chamber. Thereafter, brains were fixed, stained and imaged using the aforementioned protocols for immunohistochemistry and microscopy (see Extended data Fig. 14a).

In the presence of 20-HydroxyEcdysone in the media, strong expression of EcR-B1 and Hr3 is observed throughout the optic lobe, while no expression of Ftz-f1 is seen (Extended data Fig. 14a). When 20-HydroxyEcdysone is omitted from the culture medium, no expression of Hr3 is seen (as expected), however weak induction of Ftz-f1 is observed. The transcriptional regulation of ftz-f1 is not completely understood, and previous work has shown that ftz-f1 can be induced even without Hr3 expression. Apart from Hr3, another target of EcR – Blimp-1 has also been shown to regulate ftz-f1 expression by functioning as a repressor. Indeed, loss of Blimp-1 leads to ftz-f1 expression^17^. In brains incubated without Ecdysone, the weak induction of ftz-f1 may be (in part) due to lack of Blimp-1 expression.

### Immunohistochemistry and microscopy

Immunohistochemistry and microscopy were performed as described in Xu *et al.*^62^ with the following modifications: 1) For experiments involving staining of presynaptic sites and L5 morphology, brains were fixed using glyoxal (3.12% glyoxal, 0.75% acetic acid and 20% ethanol, pH adjusted to 5.0) for 30’ at RT and then washed 3X with PBST. 2) All images were acquired using an LSM880 confocal microscope.

Primary antibodies used in this study were: mAb24B10 (1:20, DSHB), rabbit anti-NetB (1:500, gift from Akin Lab), rabbit anti-dsRed (1:400, Clontech 632496), chicken anti-GFP (1:1000, Abcam ab13970), rat anti-Flag (1:200, Novus Biologicals, NBP1-06712), mouse anti-EcR-B1 (1:20, DSHB AD4.4), mouse anti-EcR-A (1:10, DSHB 15G1a), rabbit anti-Hr3 (1:50, gift from Thummel Lab), mouse anti-Svp (1:20, DSHB 5B11), rabbit anti-Erm (1:100, gift from Wang Lab), rat anti-Bab2 (1:500, gift from Laski Lab), mouse anti-V5 (1:200, BioRad MCA1360), guinea pig anti-Bsh (1:400, see Tan *et al.* 2015), guinea-pig anti-DIP-β (1:300, gift from Pecot Lab). Secondary antibodies used in this study were: goat anti-mouse 488 (1:500, ThermoFisher A-32723), goat anti-mouse 568 (1:500, ThermoFisher A-11031), goat anti-mouse 647 (1:500, ThermoFisher A-21235), goat anti-chicken 488 (1:1000, ThermoFisher A-11039), goat anti-rabbit 568 (1:500, ThermoFisher A-11011), goat anti-guinea pig 647 (1:500, ThermoFisher A-21450), goat anti-rabbit 647 (1:200, ThermoFisher A-32733), goat anti-rat 568 (1:500, ThermoFisher A-11077), goat anti-guinea pig 568 (1:500, A-11075).

### Image analysis

All images for figures were created using ImageJ (v2.1.0) or Imaris (v9.1.2). Details for quantification of phenotypes are given below.

#### R8 targeting

Imaris (v9.1.2) was used to measure the distance of R8 terminals from the top of the medulla (M0), as well as the distance of 6th medulla layer (M6) from the top of the medulla. The latter was estimated using mAb24B10 staining, which labels R7 neurons that terminate in M6. Values reported on the X-axis of Fig. 4d and Extended data Fig. 15b are (R8-M0)/ (M6-M0).

#### L5 morphological defects

Imaris was used to visualize individual neurons. Each neuron was then manually assigned to one of the stated categories.

#### Quantification of BRP puncta

For lamina neuron presynaptic sites in the lamina: the distance between the proximal end of the lamina and the distal end of the lamina was measured using Imaris individually for all lamina cartridges. A line was drawn at the measured distance divided by 2. All BRP puncta more distal of this line were counted manually. For R8 presynaptic sites: All Brp puncta were counted manually.

### Other statistics

One-sided Fisher’s exact test was used to determine the enrichment of dynamic genes (temporal dynamicity score > 0, calculated as in Fig. 1a, but specifically for L3) within common targets of EcR and Erm as compared to targets of Erm alone. Targets of Erm were obtained from Peng *et al.* and Santiago *et al.*^38, 39^.

For all box-plots, solid line depicts median, while the upper and lower bounds of the box depict the third and first quantile of the data spread respectively.

Kolmogorov-Smirnov (KS) test (Fig. 4d, Extended data Fig. 15b, c), two-tailed Student’s t-test (Fig. 2b-e, 2j, 4c, Extended data Fig. 6, 8c, 15a), two-tailed Fisher’s exact test (Fig. 2g, h, Extended data Fig. 5a), one-tailed Fisher’s exact test (Fig. 3b, Extended data Fig. 2d) were performed using the following basic R functions (R 3.6.1) respectively: ks.test, t.test, fisher.test. Raw data for all analyses is available upon request.

### Data and code availability

All raw sequencing data and codes will be provided upon request. They will be made publicly available prior to publication.

---

**Extended data Fig. 1.**
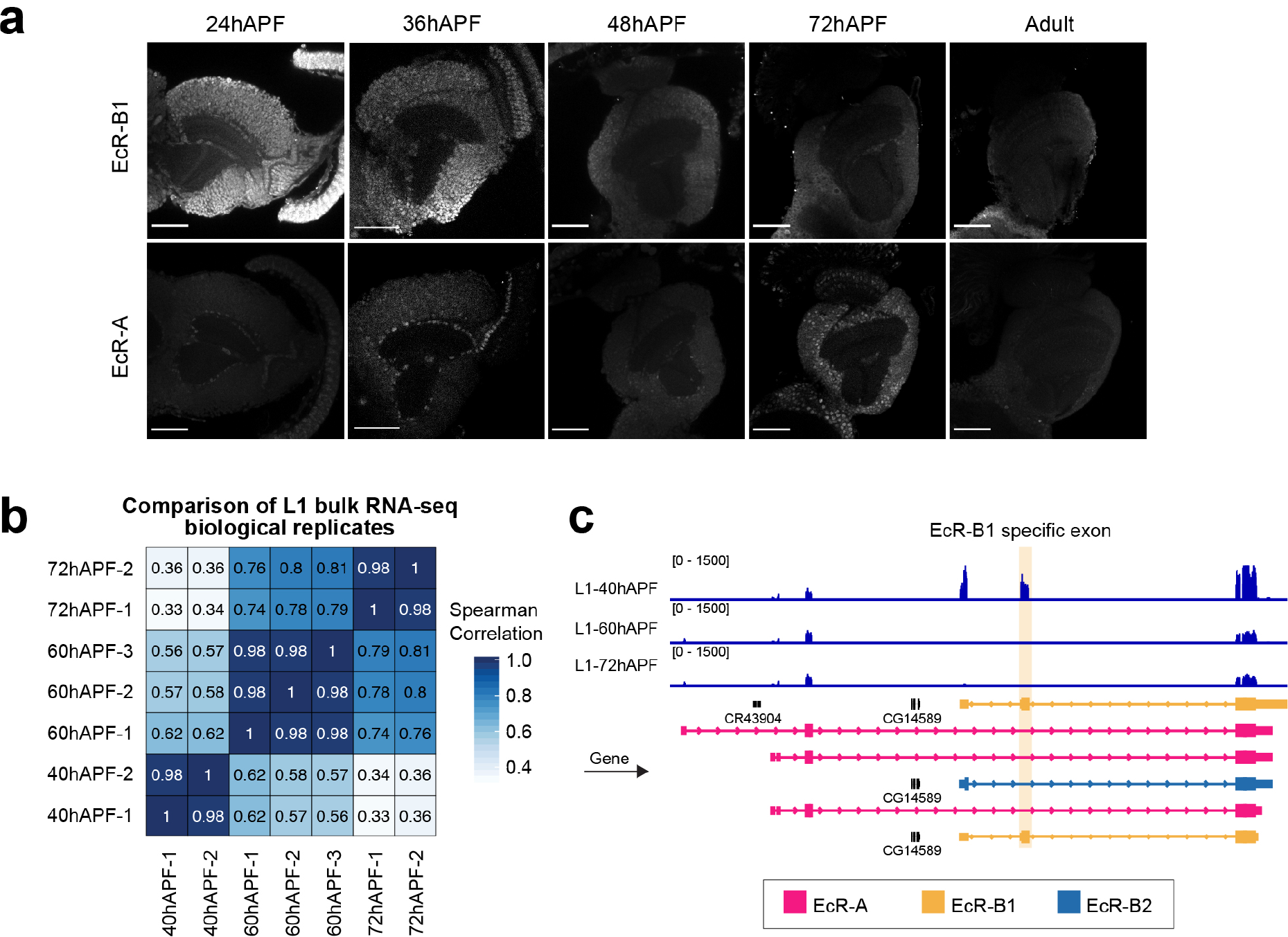
Development change from EcR-B1 to EcR-A isoform. **a,** Staining using anti-EcR-B1 or anti-EcR-A antibodies at the indicated times in development. Scale bar, 50μm. Note: EcR-A positive cells at 36hAPF are glia. **b,** Comparison of replicates of bulk RNA-Seq of L1 neurons at 40hAPF, 60hAPF and 72hAPF. Values given are Spearman correlation values (see Methods). **c,** Coverage tracks from L1 bulk RNA-Seq in the EcR locus at 40hAPF, 60hAPF and 72hAPF. EcR-A, EcR-B1 and EcR-B2 transcripts are shown, and EcR-B1 specific exon is highlighted.

**Extended data Fig. 2.**
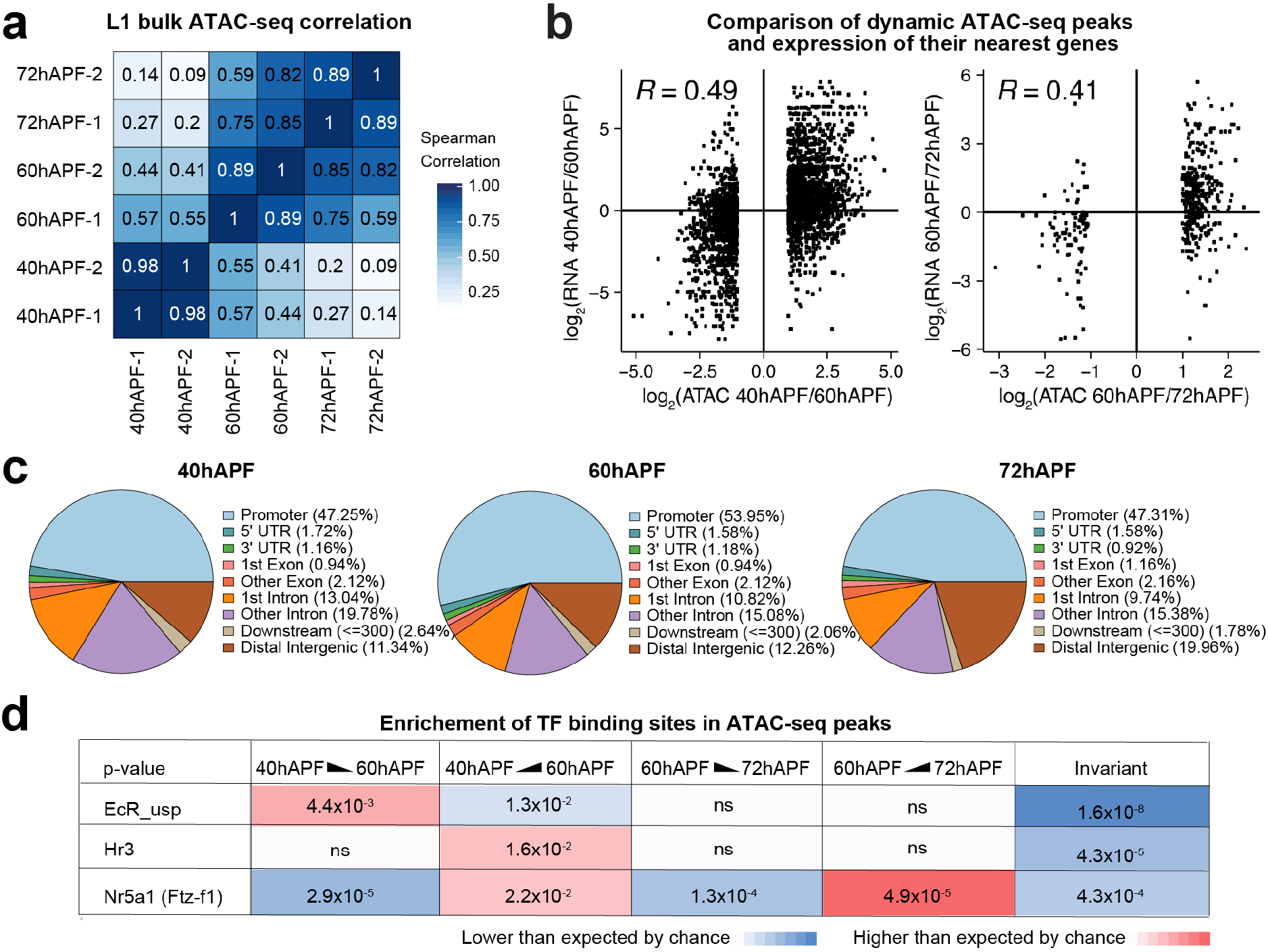
ATAC-Seq analysis of developing L1. **a,** Comparison of replicates of bulk ATAC-Seq data of L1 neurons at 40hAPF, 60hAPF and 72hAPF. Values shown are Spearman correlation values (see Methods). **b,** Comparison of change of ATAC-seq peak coverage (for regions with dynamic coverage over time) and change in expression of nearest gene. log_2_(fold change RPKM of nearest gene) vs log_2_(fold change ATAC-seq peak coverage) between 40hAPF and 60hAPF, and 60hAPF and 72hAPF. r, Pearson’s correlation coefficient. **c,** Distribution of the top 5000 peaks at each time point between various genomic landmarks. **d,** p-values (Hypergeometric test) for enrichment of binding motifs of EcR-usp complex, Hr3 and Nr5a1 (mammalian homolog for Ftz-f1) amongst the following sets of ATAC-Seq peaks: peaks going down from 40hAPF to 60hAPF, peaks going up from 40hAPF to 60hAPF, peaks going down from 60hAPF to 72hAPF, peaks going up from 60hAPF to 72hAPF, and peaks invariant over time. Red: occurrence of motif is higher than expected by chance, blue: occurrence of motif is lower than expected by chance (see Methods).

**Extended data Fig. 3.**
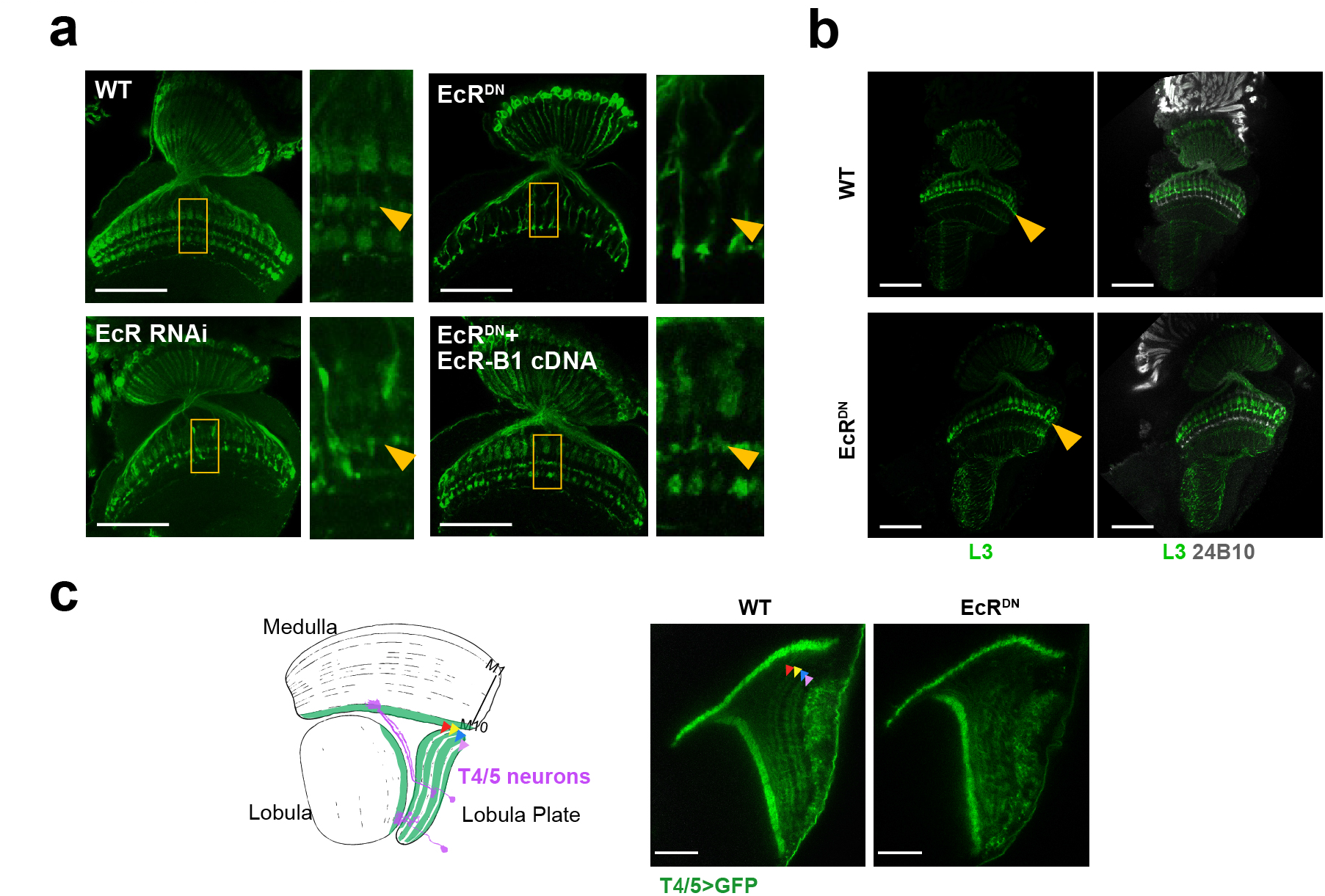
Analysis of morphology with or without EcR^DN^ expression. **a,** Morphology of lamina neurons (L1-L5) in wildtype (WT) brains and upon pan-lamina expression (using 9B08 Gal4) of EcR^DN^, EcR^DN^ + EcR-B1 cDNA or EcR RNAi. Arrowhead in inset points to M3 medulla layer. Note: loss of arborization in M3 with EcR^DN^ is likely due to loss of driver expression in L3 neurons (see Extended data Fig. 3b). **b,** Morphology of L3 neurons with or without EcR^DN^ expression using an L3-specific driver (9D03 Gal4). Note, in adults 9D03 Gal4 also labels some L2 neurons. This is not the case during development (see Extended data Fig. 16b, c) **a, b,** Arrowhead points to M3 layer in the medulla. Scale bar, 50μm. **c,** Effect of EcR^DN^ expression on morphology of T4/T5 neurons. Four layers in the lobula plate, a, b, c and d, are marked with red, yellow, blue and pink arrowheads, respectively. Cartoons of one T4 (purple neuron, top) and one T5 neuron (purple neuron, bottom) are shown to highlight wildtype morphology. Scale bar, 20µm.

**Extended data Fig. 4.**
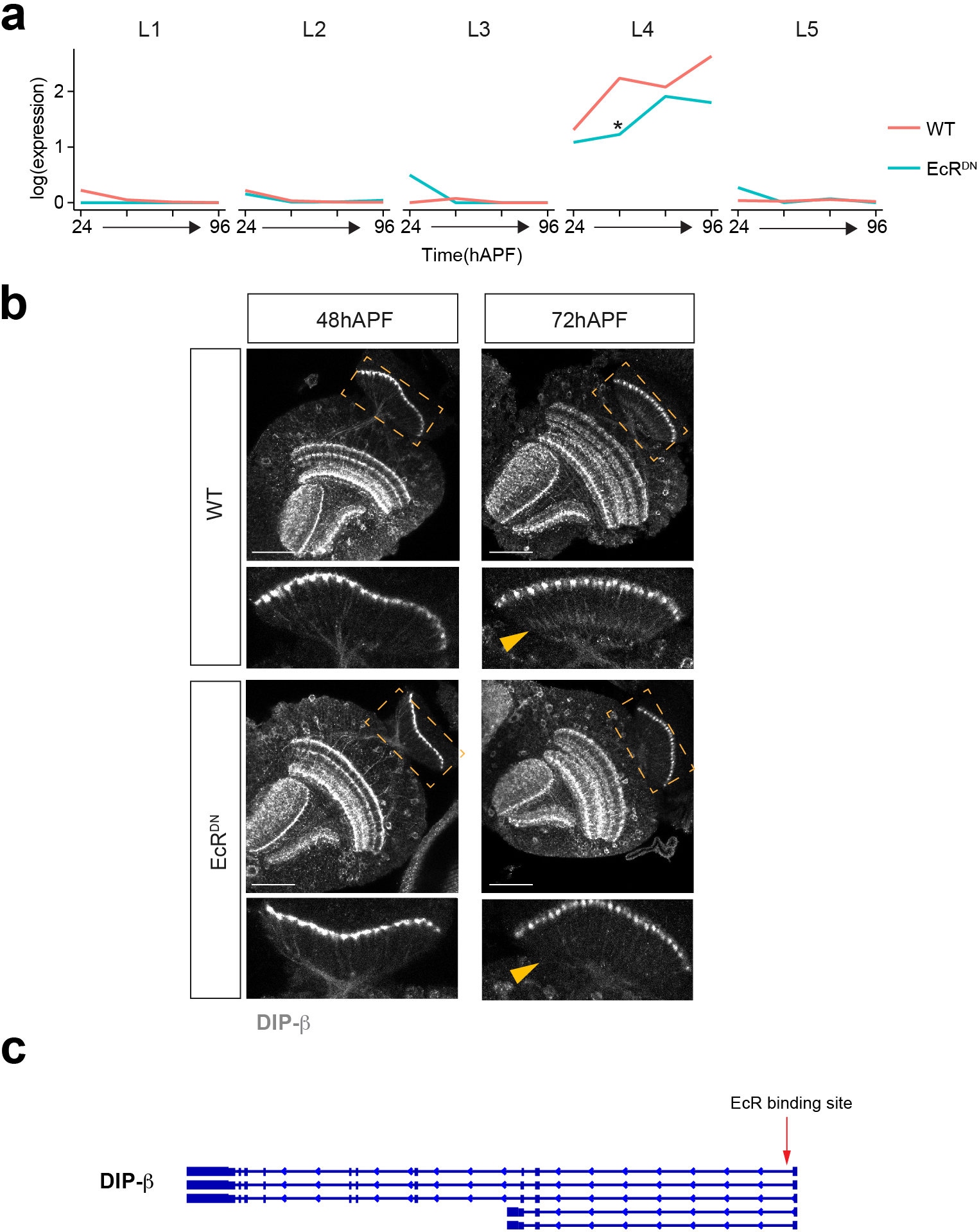
DIP-β expression with or without EcR^DN^. **a,** DIP-β mRNA expression in L1-L5 neurons across development with or without pan-lamina expression of EcR^DN^ (from scRNA-Seq based transcriptomic analyses, see Fig. 3, Extended data Fig. 7, 8). * significant difference between control and EcR^DN^ (change in expression > 2fold, p-value < 0.05). **b,** Staining using anti-DIP-β antibody at the indicated time points in development with or without pan lamina expression of EcR^DN^. Inset shows the lamina neuropil. Arrowhead points to proximal lamina neuropil, positive for DIP-β staining at 72hAPF. Note that staining is largely absent from the lamina neuropil (yellow arrowheads). The level of DIP-β RNA is reduced in EcR^DN^ at 48 but rises to near normal levels again at 72 hrs. We assume that the decrease in RNA leads to a lag in the accumulation of DIP-β protein. **c,** EcR-usp complex binding motif within the first intron of DIP-β (FDR < 10^-7^, see Methods).

**Extended data Fig. 5.**
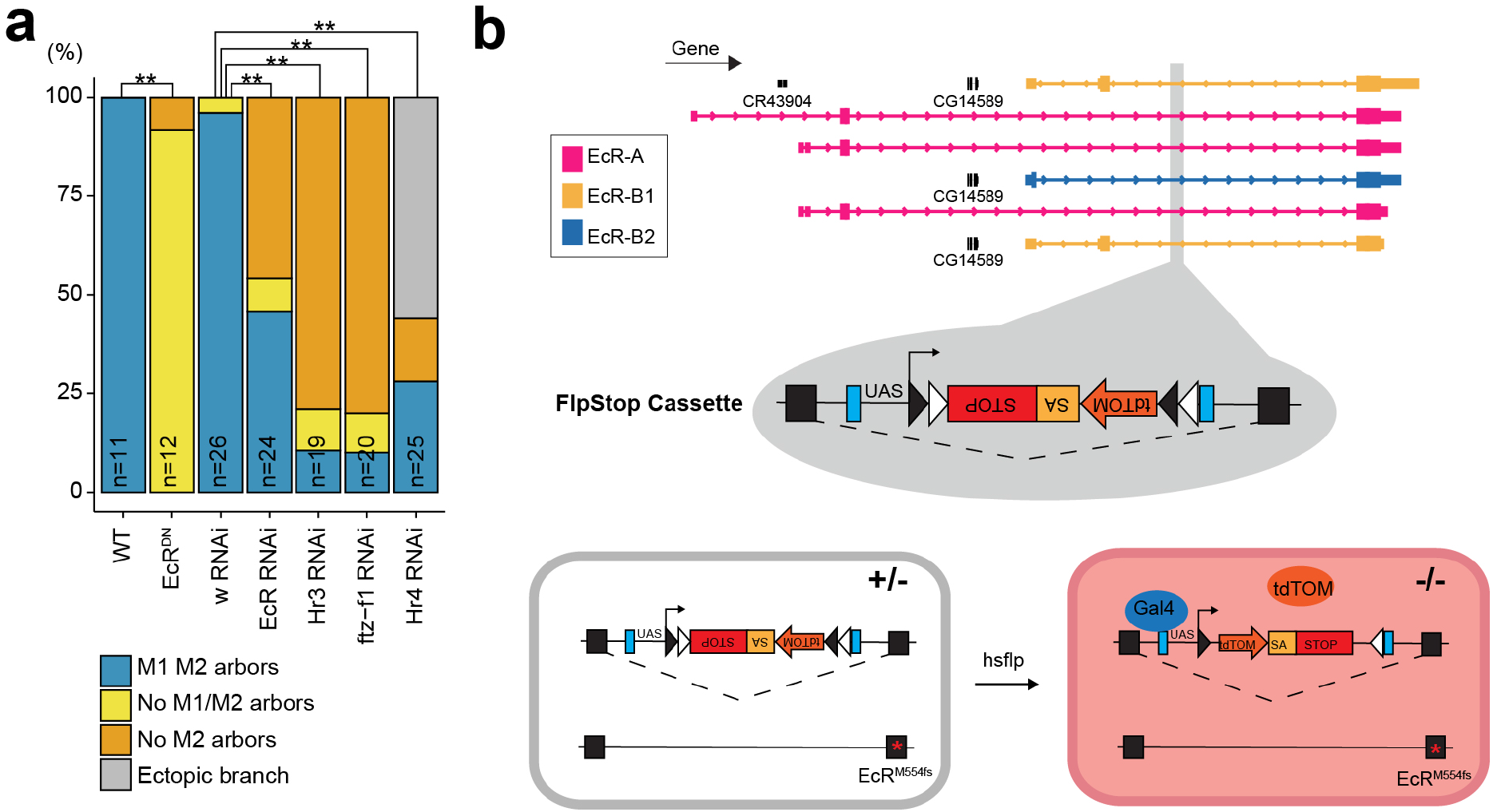
Cell and cell-type autonomous analysis of EcR function. **a,** Different L5 arborization defects and their distributions under the given genotypes (see Fig. 2g). All transgenes are expressed using an L5-specific driver (pan-lamina driver is used for data in Fig. 2g). n = number of neurons (all conditions, animals ≥ 4). **, p-value < 0.001. **b,** schematic showing the EcR^FlpStop^ allele. FlpStop cassette is inserted in the first common intron of EcR (grey bar shows insertion site, Mi{MIC}EcRMI05320). Cells expressing Gal4, within which a stochastic, heat-shock FLP recombinase-mediated flipping of the cassette occurs, express tdTom.

**Extended data Fig. 6.**
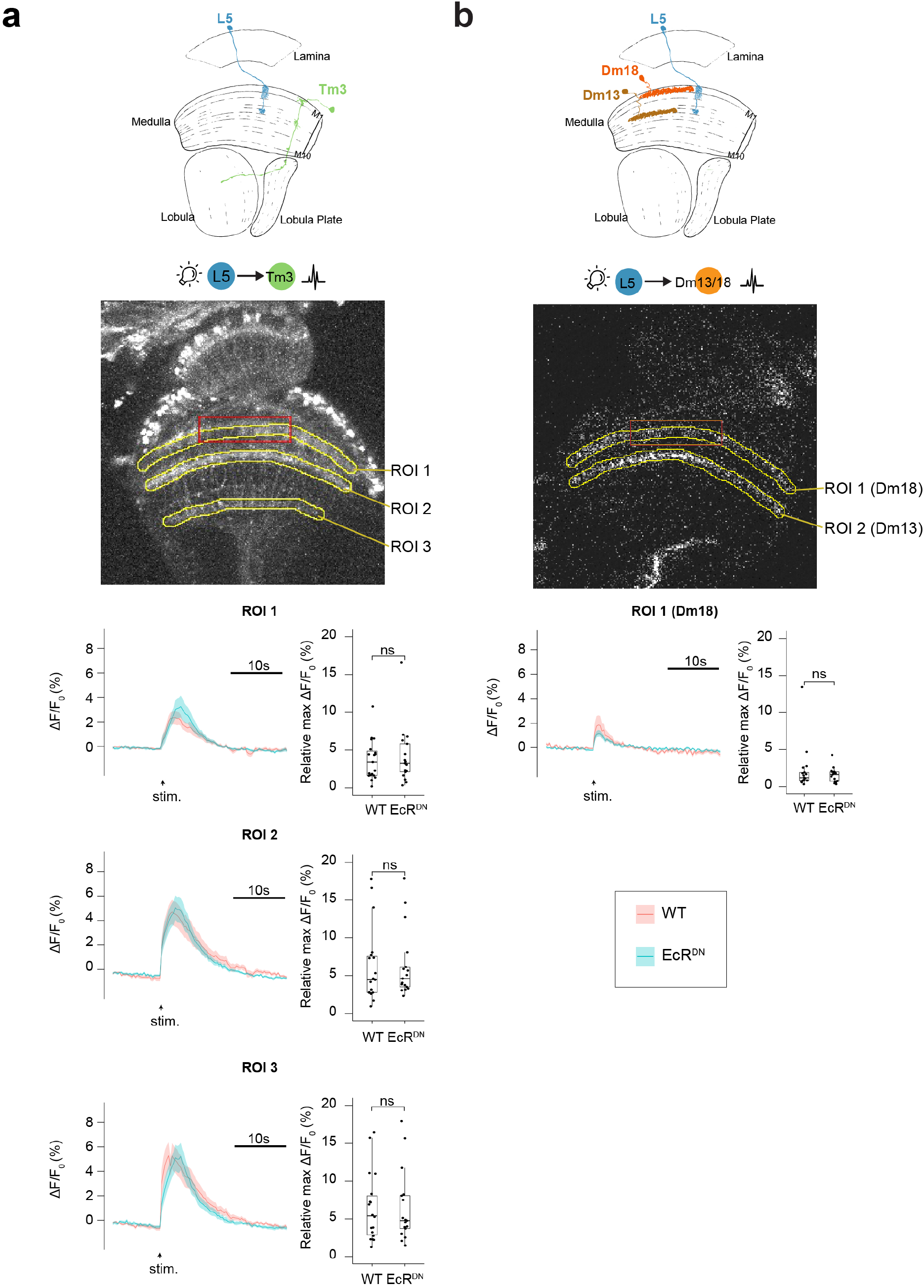
Optogenetics-based assay for communication between L5 and its postsynaptic partners. **a,** Ca^2+^ response from Tm3 (measured using GCaMP6s) upon optogenetic stimulation of L5 with (blue line) or without (red line) EcR^DN^ expression in L5. GCaMP6s responses were measured in 3 regions of interest (ROI). ROI 1, ROI 2 and ROI 3 span medulla layers M1, M5 and M10 respectively. (WT, 19 animals; EcR^DN^, 17 animals.) **b,** Ca^2+^ response from Dm13 and Dm18 (common LexA driver used yields expression in both Dm13 and Dm18, see Extended data Table 1) upon optogenetic stimulation of L5 with (blue line) or without (red line) EcR^DN^ expression in L5. ROI 1 spans M1 and measures response from Dm18. ROI 2 spans M5 and measures response from Dm13. (WT, 19 animals; EcR^DN^, 17 animals.) Note: weak response from Dm18 upon stimulation of L5 irrespective of condition. **a, b,** Amplitude of relative peak response for each condition is quantified. There is no significant difference between WT and EcR^DN^ for any comparison shown here. For Dm13 response, see Fig. 2j.

**Extended data Fig. 7.**
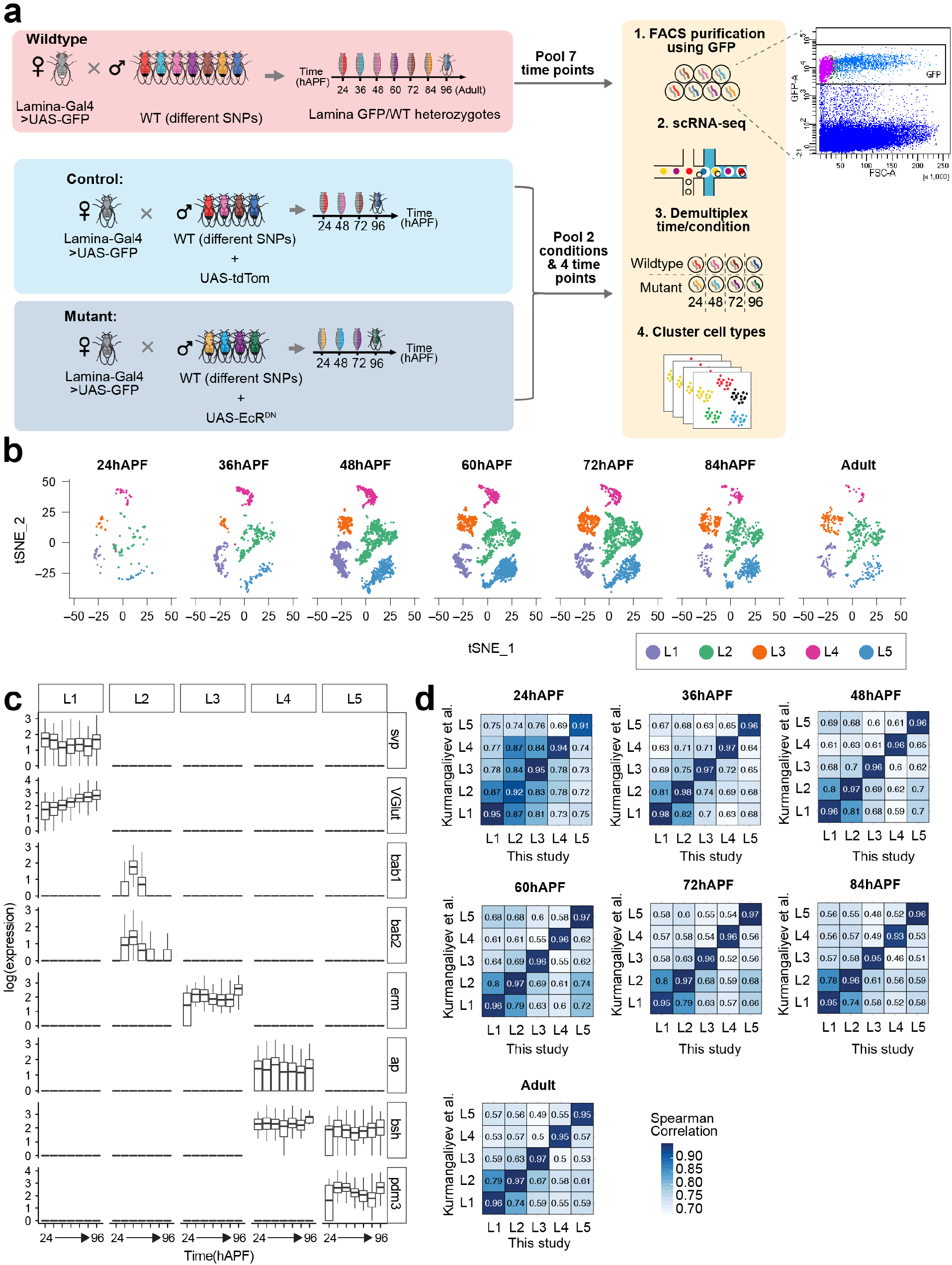
Approach for identification of EcR and Hr3 targets using scRNA-Seq. **a,** Scheme for scRNA-Seq based transcriptomic analysis of WT and mutant lamina neurons (see Kurmangaliyev *et al.*^12^ and methods). GFP vs forward scatter 2-D plot showing criteria used to enrich for lamina neurons by FACS is shown on the right. ‘Cells’ highlighted in purple were excluded despite being GFP+ due to their small size. **b,** tSNE plots showing WT L1-L5 clusters at 24, 36, 48, 60, 72, 84 and 96 hAPF (Adult). **c,** Log(expression) of previously identified lamina neuron type-specific genes in L1-L5 clusters identified at each time point over development (see Tan *et al.*^32^). **d,** Comparison of lamina neuron transcriptomes generated by scRNA-Seq in this study and by scRNA-Seq in Kurmangaliyev *et al.* Values given are Spearman correlation values.

**Extended data Fig. 8.**
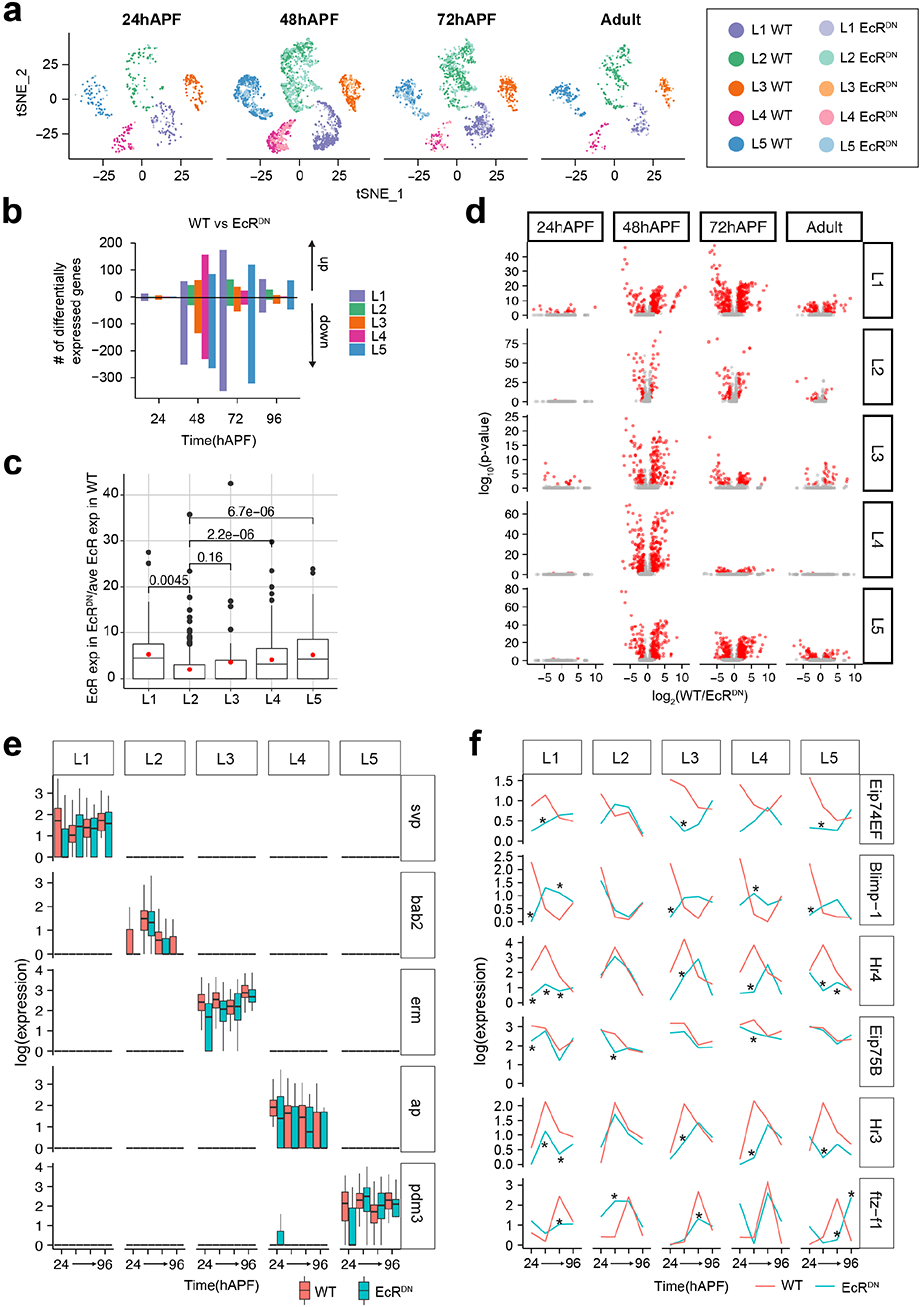
scRNA-Seq-based analysis of WT and EcR^DN^ expressing lamina neurons. **a,** tSNE plots showing WT and EcR^DN^-expressing L1-L5 clusters at 24, 48, 72 and 96 hAPF (Adult). **b**, Number of genes up or downregulated in EcR^DN^ in L1-L5 neurons. **c,** Expression of EcR in EcR^DN^-expressing lamina neurons at 48hAPF normalized to mean expression of EcR in wildtype cells at 48hAPF (done separately for each lamina neuron-type). Red dots, mean of data spread. Increase in EcR expression in EcR^DN^-expressing cells over wildtype is expected to be due to the expression of the EcR^DN^ transgene. Note the poor induction of EcR^DN^ in L2 neurons. p-value from student’s t-test are stated in the figure for comparison between L2 and other lamina neuron-types. The difference between EcR^DN^ expression in L2 and L3 neurons is not significant likely due to the low cell numbers of EcR^DN^-expressing L3 neurons. **d,** Volcano plots showing significant gene expression changes in L1-L5 neurons throughout development. Red dots: fold change > 2 and p-value < 0.05. **e,** Log(expression) of lamina neuron-type specific TFs with (blue) or without (red) EcR^DN^. Note no change in expression of TFs ± EcR^DN^. **f,** Log(expression) of TFs in the Ecdysone-pathway in WT (red lines) and EcR^DN^-expressing (blue lines) L1-L5 neurons. *, p-value < 0.05, fold change > 2.

**Extended data Fig. 9.**
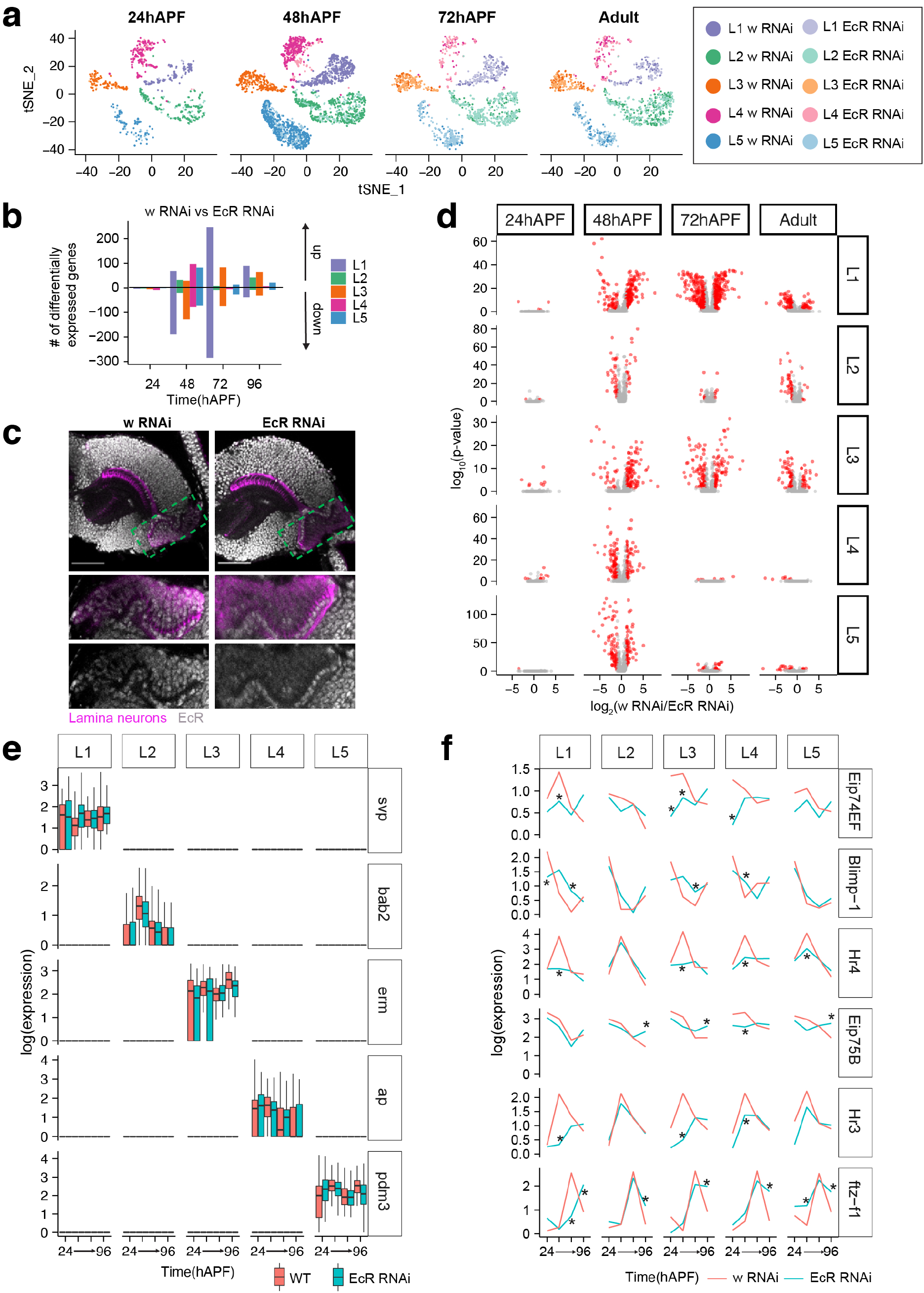
scRNA-Seq-based analysis of w RNAi and EcR RNAi expressing lamina neurons. **a,** tSNE plots showing w RNAi and EcR RNAi-expressing L1-L5 clusters at 24, 48, 72 and 96 hAPF (Adult). **b**, Number of genes up or downregulated in EcR RNAi in L1-L5 neurons. **c,** Image showing optic lobe (top) stained using an antibody targeting all EcR isoforms (grey) at 24hAPF. Box with green dotted outline marks the region containing lamina neuron cell-bodies. This region is magnified in bottom two panels. Lamina neurons are labeled in magenta. Scale bar, 50µm. **d,** Volcano plots showing significant gene expression changes in L1-L5 neurons throughout development. Red dots: fold change > 2 and p-value < 0.05. **e,** Log(expression) of lamina neuron-type specific TFs with (blue) or without (red) EcR RNAi. Note no change in expression of TFs ± EcR RNAi. **f,** Log(expression) of TFs in the Ecdysone-pathway in WT (red lines) and EcR RNAi-expressing (blue lines) L1-L5 neurons. *, p-value < 0.05, fold change > 2.

**Extended data Fig. 10.**
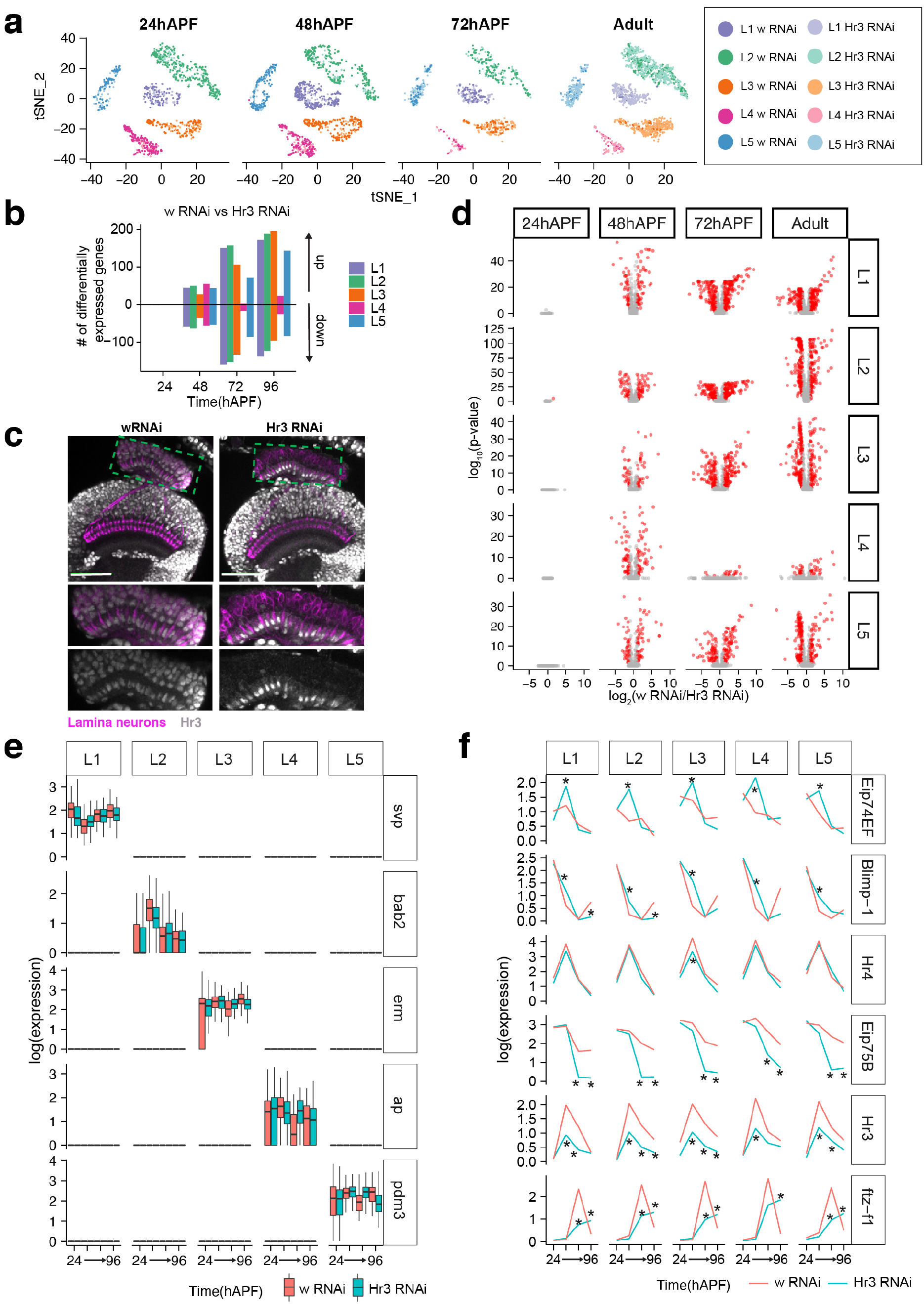
scRNA-Seq-based analysis of w RNAi and Hr3 RNAi expressing lamina neurons. **a,** tSNE plots showing w RNAi and Hr3 RNAi-expressing L1-L5 clusters at 24, 48, 72 and 96 hAPF (Adult). **b**, Number of genes up or downregulated in Hr3 RNAi in L1-L5 neurons. **c,** Image showing optic lobe (top) stained using an antibody targeting Hr3 (grey) at 24hAPF. Box with green dotted outline marks the region containing lamina neuron cell-bodies. This region is magnified in bottom two panels. Lamina neurons are labeled in magenta. Scale bar, 50µm. **d,** Volcano plots showing significant gene expression changes in L1-L5 neurons throughout development. Red dots: fold change > 2 and p-value < 0.05. **e,** Log(expression) of lamina neuron-type specific TFs with (blue) or without (red) Hr3 RNAi. Note no change in expression of TFs ± Hr3 RNAi. **f,** Log(expression) of TFs in the Ecdysone-pathway in WT (red lines) and Hr3 RNAi-expressing (blue lines) L1-L5 neurons. *, p-value < 0.05, fold change > 2.

**Extended data Fig. 11.**
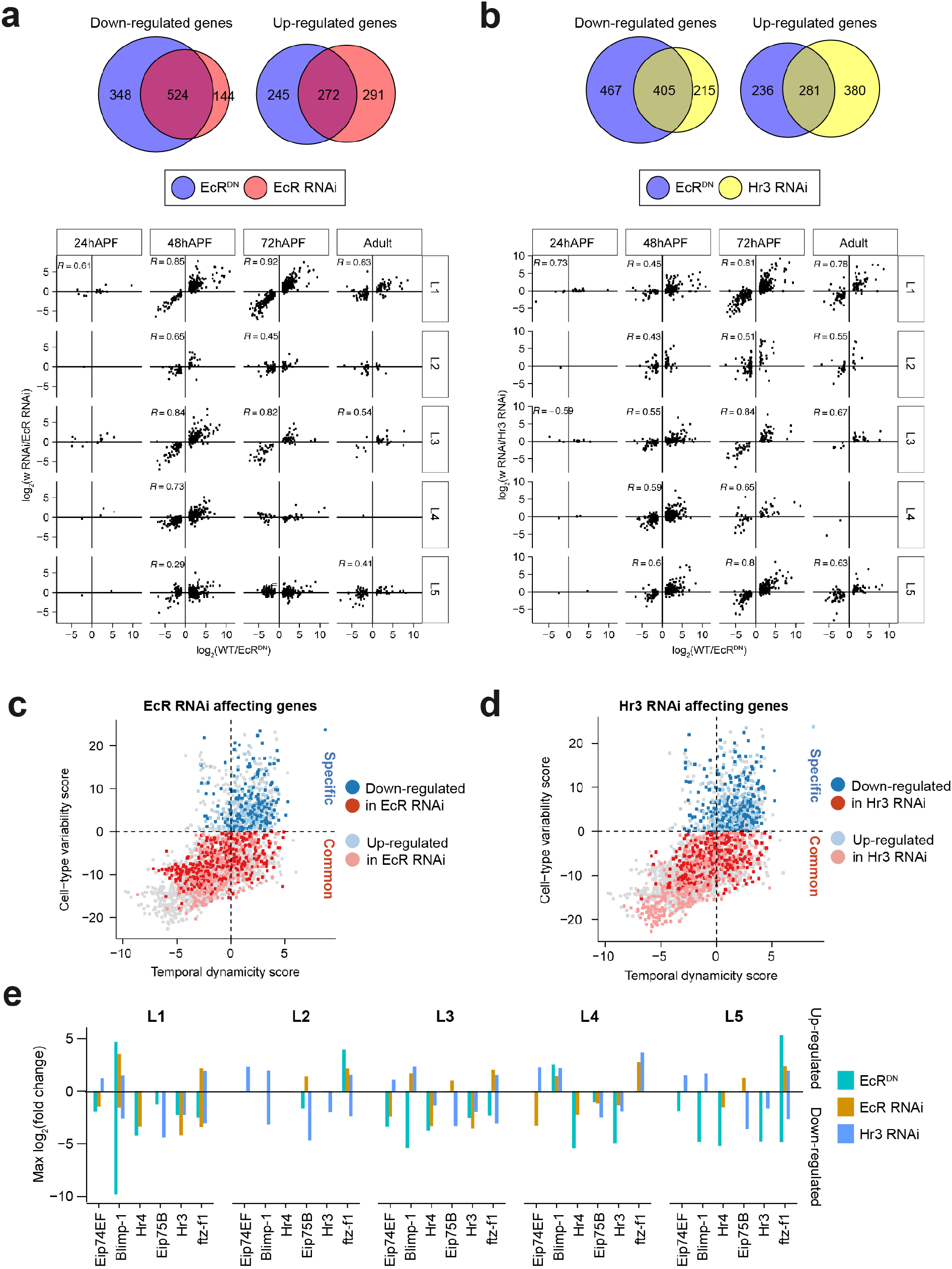
Comparison of genes affected by EcR^DN^, EcR RNAi and Hr3 RNAi. **a,** Top, Venn diagram showing overlap between genes downregulated by EcR^DN^ and EcR RNAi across all time points and lamina neuron-types. Below, log_2_(normalized expression in w RNAi/EcR RNAi) vs log_2_(normalized expression in WT/EcR^DN^) for L1-L5 neurons throughout development. Correlation coefficient, R, is given for comparisons where p < 0.05. **b,** Top, Venn diagram showing overlap between genes downregulated by EcR^DN^ and Hr3 RNAi across all time points and lamina neuron-types. Below, log_2_(normalized expression in w RNAi/Hr3 RNAi) vs log_2_(normalized expression in WT/EcR^DN^) for L1-L5 neurons throughout development. Correlation coefficient, R, is given for comparisons where p < 0.05. **c,** Cell-type variability vs Temporal dynamicity plot for EcR RNAi-affected genes (fold change > 2, p-value < 0.05). Cell-type Specific (blue) and Common (red) targets are shown. Darker colors: genes reduced in EcR RNAi, lighter colors: genes increased in EcR RNAi. **d,** Cell-type variability vs Temporal dynamicity plot for Hr3 RNAi-affected genes (fold change > 2, p-value < 0.05). Cell-type Specific (blue) and Common (red) targets are shown. Darker colors: genes reduced in Hr3 RNAi, lighter colors: genes increased in Hr3 RNAi. **e,** Maximum change in expression of Ecdysone-pathway TFs in L1-L5 neurons with EcR^DN^, EcR RNAi and Hr3 RNAi. Note that EcR RNAi often has weaker effect on TF expression as compared to EcR^DN^.

**Extended data Fig. 12.**
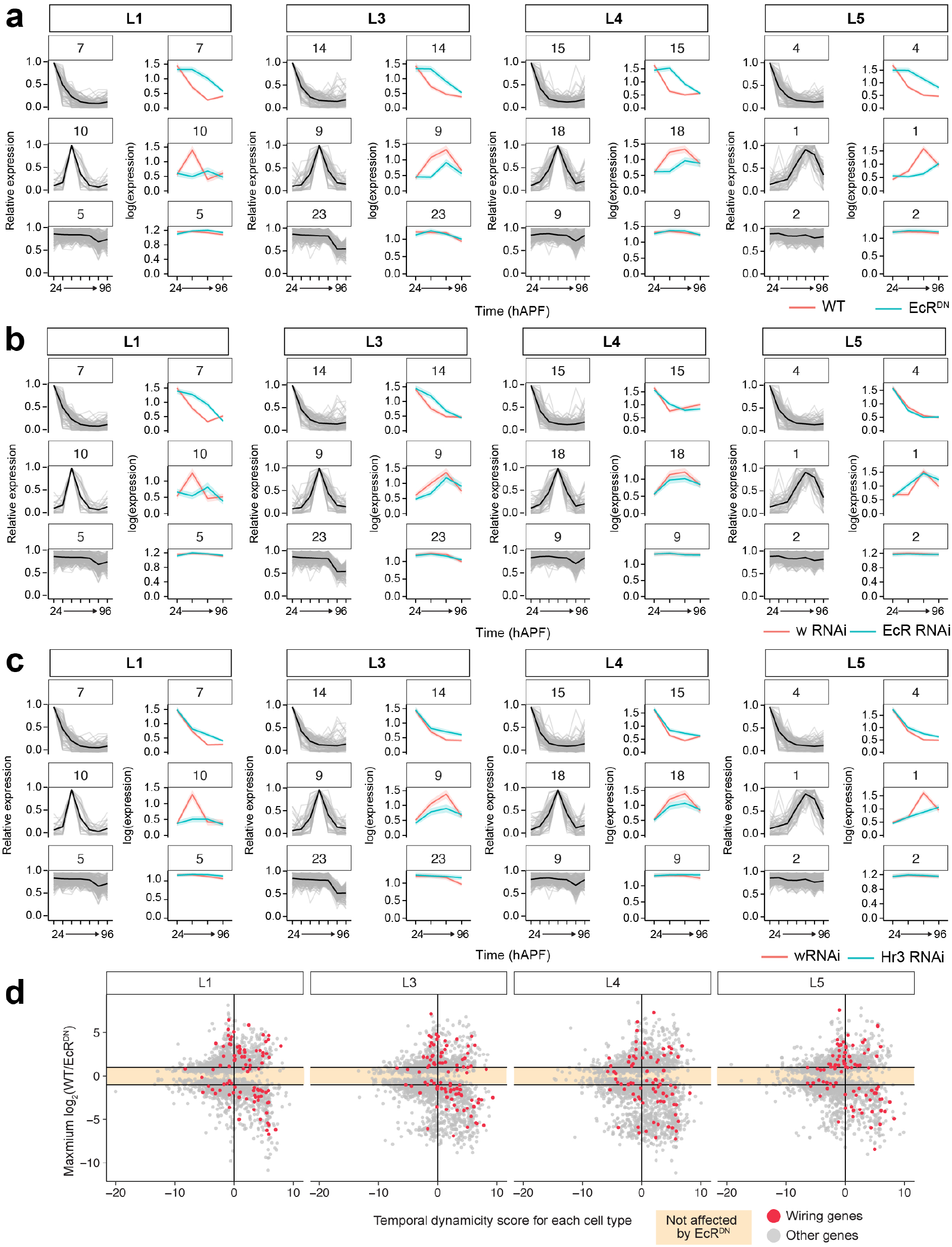
Clusters of genes most affected by EcR^DN^, EcR RNAi and Hr3 RNAi. All genes expressed in L1-L5 neurons (done separately for each cell-type) were clustered into groups (using k-means clustering) based on their expression dynamics (see Methods). Clusters that show maximum upregulation or downregulation with EcR^DN^ are shown in **a – c** (clusters are indicated in numbers above each graph). Also shown are clusters unchanged by EcR^DN^. For each panel: left, light grey lines, relative expression of all genes in the cluster; black line, mean of relative expression of all genes in the cluster. Right, red line, mean relative expression in control; blue line, mean relative expression with perturbation. Shades are SEM. d, examples of dynamic wiring genes that are not affected by EcR^DN^. **d,** Plots showing maximum change in expression [log_2_(WT/EcR^DN^)] caused by EcR^DN^ vs the temporal dynamicity of the gene (calculated separately for each cell-type). Shown in red are wiring genes. Colored region on the plot represents genes not affected by EcR^DN^. Note many genes including wiring genes (especially in L1, L3 and L4) with high temporal dynamicity scores that are not affected by EcR^DN^.

**Extended data Fig. 13.**
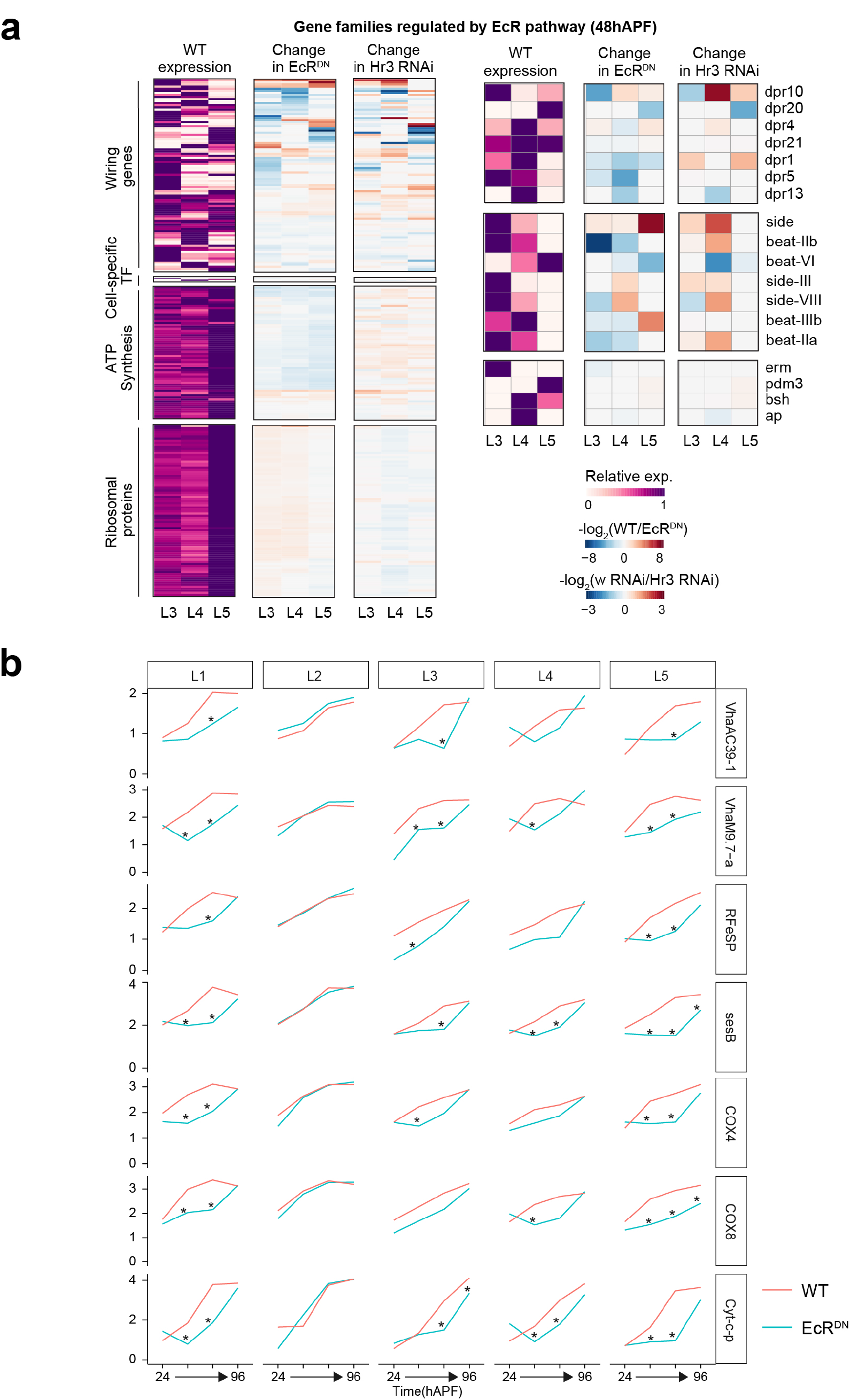
Families of genes affected by EcR^DN^ and Hr3 RNAi. **a,** Relative WT expression (left), change in expression with EcR^DN^ (center), and change in expression with Hr3 RNAi (right) shown as heat maps for all genes expressed in L3-L5 neurons belonging to the specified gene categories at 48hAPF (also see Extended data Fig. 13). Examples of genes belonging to the Dpr family, Side-Beat family and cell-type specific transcription factors are shown separately. **b,** Expression of some genes involved in ATP synthesis and vacuolar ATPase biology are shown with and without EcR^DN^ in L1-L5 neurons across development. *, fold change between WT and EcR^DN^ > 2, p-value < 0.05. Note: expression in adults is not significantly affected by EcR^DN^.

**Extended data Fig. 14.**
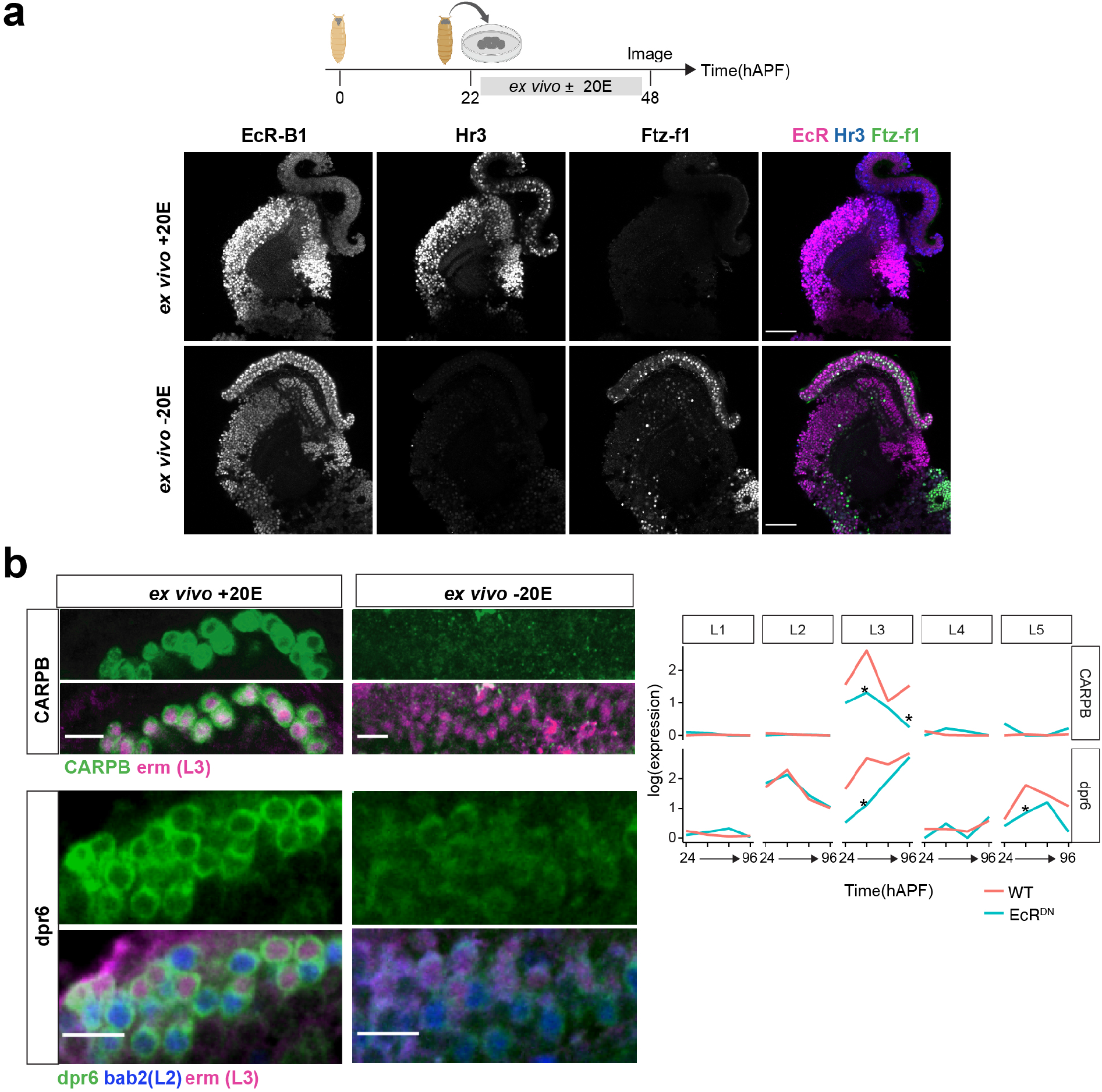
*ex vivo* culture of pupal brain with or without Ecdysone. **a,** Top, schematic of experimental setup (see Methods). Briefly, brains are dissected at 22hAPF then incubated for 26h in media ± 20E (20 HydroxyEcdysone, active form of Ecdysone). Bottom, optic lobes stained for EcR-B1, Hr3 and Ftz-f1 cultured *ex vivo* ± 20E. Scale bar, 50μm. **b,** Left, Staining for CARPB and dpr6 reporters (MiMIC lines, see Methods) in lamina neuron cell-bodies ± 20E. Right, CARPB and dpr6 expression with (blue) or without (red) EcR^DN^. *, fold change between WT and EcR^DN^ > 2, p-value < 0.05. Note: dpr6 expression is unchanged in L2 with pan-lamina expression of EcR^DN^, however staining in brains cultured *ex vivo* without 20E in the medium shows reduced protein expression in L2 (bab2 positive cell-bodies). This is consistent with weak activity of the pan-lamina driver (9B08 Gal4) in L2 at 48hAPF leading to ineffective inhibition of the Ecdysone-pathway (see Extended data Fig. 8c).

**Extended data Fig. 15.**
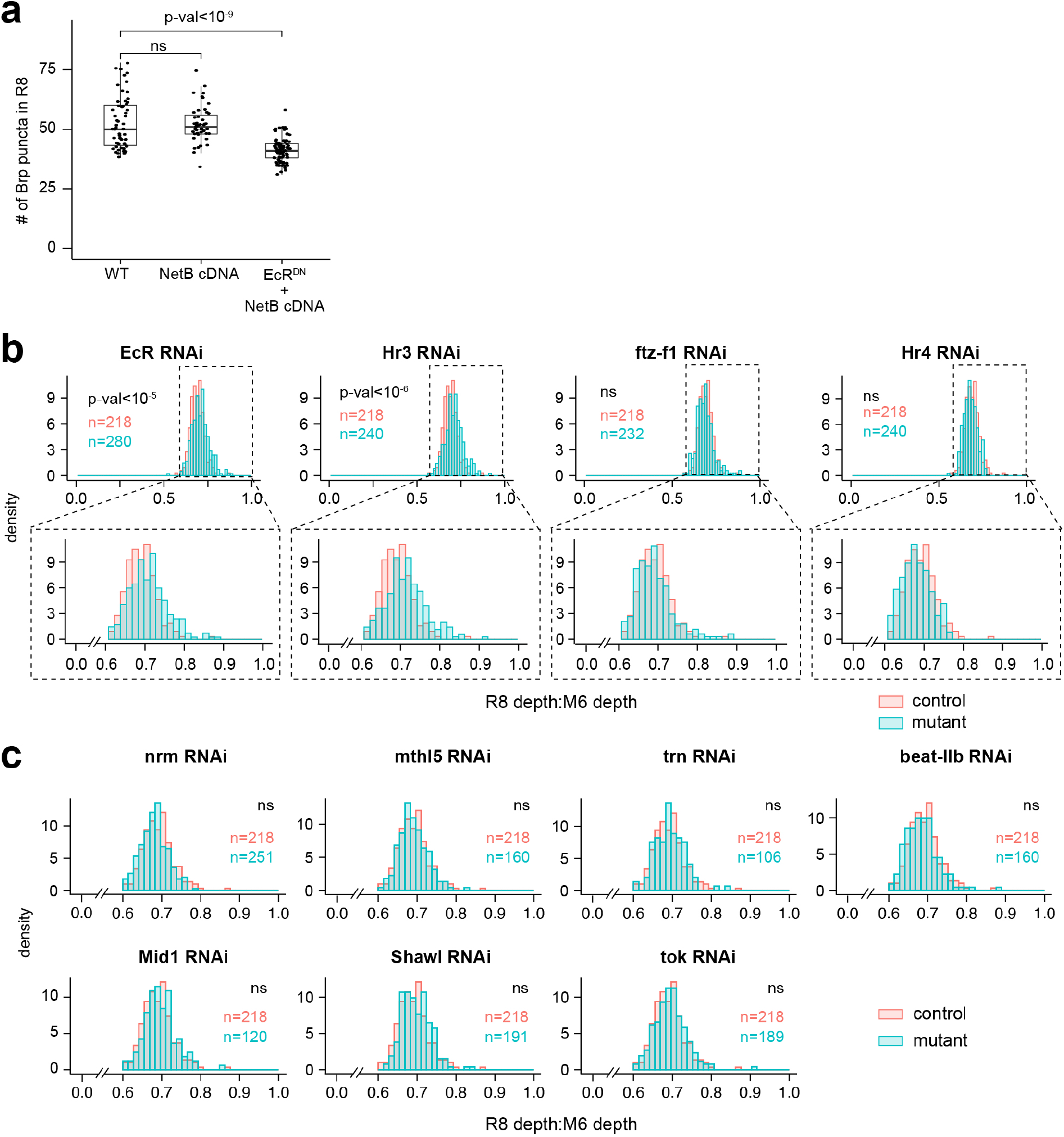
Genes expressed in L3 required for R8 wiring. **a,** Total number of presynaptic sites (Brp puncta) in R8 neurons with WT, NetB overexpressing, or EcR^DN^ and NetB-expressing L3 neurons. NetB overexpression is unable to rescue the reduction in R8 presynaptic sites seen with expression of EcR^DN^ in L3 neurons (see Fig. 4c). p-value (Student’s t-test) is given. **b, c,** Distributions of R8 axon terminal depth in w RNAi (red) or with expression of other RNAi (as shown, blue) in L3 neurons. All conditions, number of animals ≥ 3. p-values (Kolmogorov-Smirnov test) are given. ns = Not significant. **b,** RNAi against TFs in the Ecdysone-pathway. **c,** Results from RNAi screen showing genes that did not significantly affect R8 axon depth. Numbers of neurons/ condition are given.

**Extended data Fig. 16.**
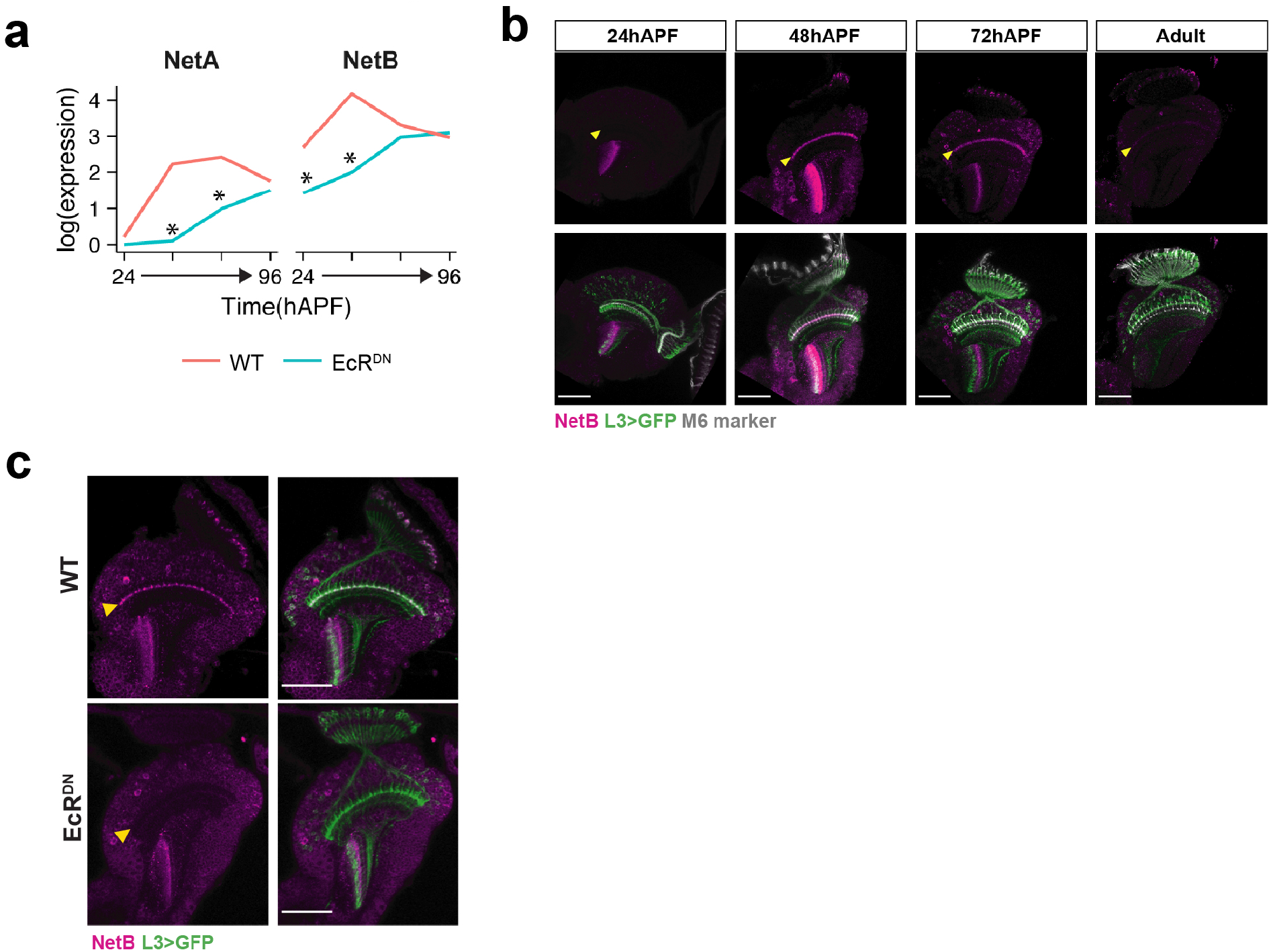
Netrin expression in L3 requires EcR activity. **a,** Expression of NetA and NetB in L3 with (blue) or without (red) EcR^DN^ expression. *, fold change between WT and EcR^DN^ > 2, p-value < 0.05. **b,** Staining using anti-NetB antibody (magenta) at the indicated times in development. Marker for M6 medulla layer (24B10, grey) and L3 neurons labeled with GFP (green) are shown. **c,** Staining using anti-NetB antibody (magenta) ± EcR^DN^ expression only in L3. L3 neurons labeled with GFP (green) are shown. **b, c,** Scale bar, 50μm. Yellow arrowhead, M3 medulla layer.

## Extended data Tables

**Extended data Table 1.** Fly strains used in this study

**Extended data Table 2.** Temporal dynamicity and cell-type variability scores

**Extended data Table 3.** Wiring genes

**Extended data Table 4.** Gene ontology and Reactome analyses

**Extended data Table 5.** L1 bulk ATAC-Seq

**Extended data Table 6.** scRNA-Seq data for wildtype Lamina neurons

**Extended data Table 7.** scRNA-Seq data for wildtype vs EcR^DN^-expressing Lamina neurons

**Extended data Table 8.** scRNA-Seq data for wildtype vs EcR RNAi-expressing Lamina neurons

**Extended data Table 9.** scRNA-Seq data for wildtype vs Hr3 RNAi-expressing Lamina neurons

## Acknowledgements

We thank Dr. Pecot (Harvard), Dr. Yamanaka (UC Riverside), Dr. Black (UCLA), Dr. De Robertis (UCLA), Dr. Riddiford (Univ. of Washington) and Dr. Truman (Janelia Research Campus) for helpful discussions. We thank Dr. Schuldiner (Weizmann Institute of Science) and members of the Zipursky lab for feedback on the manuscript, and Dr. Diaz de la Loza for help with figure illustrations. We would like to specifically acknowledge Juyoun Yoo’s (Zipursky Lab) and Rachel Hodge’s (Jones Lab, UCLA) help with ATAC-Seq library prep and immunostaining. We also thank the BSCRC Sequencing Core (UCLA) and the TCGB core (UCLA) for help with library preparation and sequencing; the BSCRC FACS core (UCLA) and the Witte lab (UCLA) for assistance with FACS purification of lamina neurons; and the IDRE Statistics Consulting (UCLA) and Dr. Balliu (UCLA) for assistance with the statistical analysis of data. Reagents provided by Dr. Akin (UCLA), Dr. Laski (UCLA), Dr. Wang (Duke-NUS), Dr. Pecot (Harvard), Dr. Thummel (Univ. of Utah) and Dr. Bashaw (Univ. of Penn.), and fly lines from the Bloomington Drosophila Stock Center were critical for this work. This work was supported by NIH T32-NS048004 Neurobehavioral Genetics Training Grant (S.J.), Helen Hay Whitney Foundation (S.J.) and Whitcome Fellowship (Y.L.). S.L.Z is an investigator of the Howard Hughes Medical Institute.

## Author contributions

S.J., Y.L., Y.Z.K., J.V-A and S.L.Z. designed experiments. S.J., Y.L., P.M., S.A.L, J.V-A and B.P. acquired data. S.J., Y.L., J.V-A and Y.K. analyzed the data. S.J., Y.L. and S.L.Z. wrote the manuscript with input from all co-authors.

## Additional information

Supplementary Information is available for this paper. Correspondence and requests for materials should be addressed to: Dr. S. Lawrence Zipursky -.

## REFERENCES

1. Südhof, T. C. Towards an Understanding of Synapse Formation. Neuron 100, 276–293 (2018).

2. Zipursky, S. L. & Sanes, J. R. Chemoaffinity Revisited: Dscams, Protocadherins, and Neural Circuit Assembly. Cell 143, 343–353 (2010).

3. Hassan, B. A. & Hiesinger, P. R. Beyond Molecular Codes: Simple Rules to Wire Complex Brains. Cell 163, 285–291 (2015).

4. Yogev, S. & Shen, K. Cellular and Molecular Mechanisms of Synaptic Specificity. Annu Rev Cell Dev Bi 30, 417–437 (2014).

5. Li, H. et al. Classifying Drosophila Olfactory Projection Neuron Subtypes by Single-Cell RNA Sequencing. Cell 171, 1206–1220.e22 (2017).

6. Özel, M. N. et al. Neuronal diversity and convergence in a visual system developmental atlas. Nature 1–8 (2020) doi:10.1038/s41586-020-2879-3.

7. Hobert, O. Chapter Twenty-Five - Terminal Selectors of Neuronal Identity. in Current Topics in Developmental Biology (ed. Wassarman, P. M.) vol. 116 455–475 (Academic Press, 2016).

8. Hong, W. & Luo, L. Genetic Control of Wiring Specificity in the Fly Olfactory System. Genetics 196, 17–29 (2014).

9. Dasen, J. S. & Jessell, T. M. Chapter Six Hox Networks and the Origins of Motor Neuron Diversity. in Current Topics in Developmental Biology vol. 88 169–200 (Academic Press, 2009).

10. Larkin, A. et al. FlyBase: updates to the Drosophila melanogaster knowledge base. Nucleic Acids Res 49, D899–D907 (2020).

12. Kurmangaliyev, Y. Z., Yoo, J., Valdes-Aleman, J., Sanfilippo, P. & Zipursky, S. L. Transcriptional Programs of Circuit Assembly in the Drosophila Visual System. Neuron (2020) doi:10.1016/j.neuron.2020.10.006.

11. Scheffer, L. K. et al. A connectome and analysis of the adult Drosophila central brain. Elife 9, e57443 (2020).

13. Reilly, M. B., Cros, C., Varol, E., Yemini, E. & Hobert, O. Unique homeobox codes delineate all the neuron classes of C. elegans. Nature 584, 595–601 (2020).

14. Truman, J. W., Talbot, W. S., Fahrbach, S. E. & Hogness, D. S. Ecdysone receptor expression in the CNS correlates with stage-specific responses to ecdysteroids during Drosophila and Manduca development. Dev Camb Engl 120, 219–234 (1994).

15. Riddiford, L. M., Cherbas, P. & Truman, J. W. Ecdysone receptors and their biological actions. Vitamins Hormones 60, 1–73 (2000).

16. White, K. P., Hurban, P., Watanabe, T. & Hogness, D. S. Coordination of Drosophila metamorphosis by two ecdysone-induced nuclear receptors. Sci New York N Y 276, 114–117 (1997).

17. Agawa, Y. et al. Drosophila Blimp-1 Is a Transient Transcriptional Repressor That Controls Timing of the Ecdysone-Induced Developmental Pathway. Mol Cell Biol 27, 8739–8747 (2007).

18. Rabinovich, D., Yaniv, S. P., Alyagor, I. & Schuldiner, O. Nitric Oxide as a Switching Mechanism between Axon Degeneration and Regrowth during Developmental Remodeling. Cell 164, 170–182 (2016).

19. Pak, M. D. & Gilbert, L. I. A Developmental Analysis of Ecdysteroids During the Metamorphosis of extitDrosophila Melanogaster. J Liq Chromatogr 10, 2591–2611 (1987).

20. Buenrostro, J. D., Wu, B., Chang, H. Y. & Greenleaf, W. J. ATAC-seq: A Method for Assaying Chromatin Accessibility Genome-Wide. Curr Protoc Mol Biology Ed Frederick M Ausubel, Et Al 109, 21.29.1–9 (2015).

21. Imrichová, H., Hulselmans, G., Atak, Z. K., Potier, D. & Aerts, S. i-cisTarget 2015 update: generalized cis-regulatory enrichment analysis in human, mouse and fly. Nucleic Acids Res 43, W57–W64 (2015).

22. Shlyueva, D. et al. Hormone-responsive enhancer-activity maps reveal predictive motifs, indirect repression, and targeting of closed chromatin. Mol Cell 54, 180–192 (2014).

23. Cherbas, L., Hu, X., Zhimulev, I., Belyaeva, E. & Cherbas, P. EcR isoforms in Drosophila: testing tissue-specific requirements by targeted blockade and rescue. Development 130, 271–284 (2003).

24. Xu, C. et al. Control of Synaptic Specificity by Establishing a Relative Preference for Synaptic Partners. Neuron 103, 865–877.e7 (2019).

25. Nern, A., Zhu, Y. & Zipursky, S. L. Local N-Cadherin Interactions Mediate Distinct Steps in the Targeting of Lamina Neurons. Neuron 58, 34–41 (2008).

26. Pecot, M. Y. et al. Sequential Axon-Derived Signals Couple Target Survival and Layer Specificity in the Drosophila Visual System. Neuron 82, 320–333 (2014).

27. Fisher, Y. E. et al. FlpStop, a tool for conditional gene control in Drosophila. Elife 6, (2017).

28. Yao, T.-P., Segraves, W. A., Oro, A. E., McKeown, M. & Evans, R. M. Drosophila ultraspiracle modulates ecdysone receptor function via heterodimer formation. Cell 71, 63–72 (1992).

29. Yao, T. P. et al. Functional ecdysone receptor is the product of EcR and Ultraspiracle genes. Nature 366, 476–479 (1993).

30. Schwabe, T., Borycz, J. A., Meinertzhagen, I. A. & Clandinin, T. R. Differential Adhesion Determines the Organization of Synaptic Fascicles in the Drosophila Visual System. Curr Biol 24, 1304–1313 (2014).

31. Takemura, S. et al. Synaptic circuits and their variations within different columns in the visual system of Drosophila. Proc National Acad Sci 112, 13711–13716 (2015).

32. Tan, L. et al. Ig Superfamily Ligand and Receptor Pairs Expressed in Synaptic Partners in Drosophila. Cell 163, 1756–1769 (2015).

33. Lee, C. W. & Peng, H. B. The Function of Mitochondria in Presynaptic Development at the Neuromuscular Junction. Mol Biol Cell 19, 150–158 (2008).

34. Zhou, H. & Liu, G. Regulation of density of functional presynaptic terminals by local energy supply. Mol Brain 8, 42 (2015).

35. Rangaraju, V., Lauterbach, M. & Schuman, E. M. Spatially Stable Mitochondrial Compartments Fuel Local Translation during Plasticity. Cell 176, 73–84.e15 (2019).

36. Gowrisankaran, S. & Milosevic, I. Regulation of synaptic vesicle acidification at the neuronal synapse. Iubmb Life 72, 568–576 (2020).

37. Özel, M. N., Langen, M., Hassan, B. A. & Hiesinger, P. R. Filopodial dynamics and growth cone stabilization in Drosophila visual circuit development. Elife 4, e10721 (2015).

38. Peng, J. et al. Drosophila Fezf coordinates laminar-specific connectivity through cell-intrinsic and cell-extrinsic mechanisms. Elife 7, (2018).

39. Santiago, I. J. et al. Drosophila Fezf functions as a transcriptional repressor to direct layer-specific synaptic connectivity in the fly visual system. Proc National Acad Sci 118, e2025530118 (2021).

40. Akin, O. & Zipursky, S. L. Frazzled promotes growth cone attachment at the source of a Netrin gradient in the Drosophila visual system. Elife 5, e20762 (2016).

41. Moffatt, N. S. C., Bruinsma, E., Uhl, C., Obermann, W. M. J. & Toft, D. Role of the Cochaperone Tpr2 in Hsp90 Chaperoning. Biochemistry-us 47, 8203–8213 (2008).

42. Brychzy, A. et al. Cofactor Tpr2 combines two TPR domains and a J domain to regulate the Hsp70/Hsp90 chaperone system. Embo J 22, 3613–3623 (2003).

43. Klucken, J. et al. ABCG1 (ABC8), the human homolog of the Drosophila white gene, is a regulator of macrophage cholesterol and phospholipid transport. Proc National Acad Sci 97, 817–822 (2000).

44. Suzuki, M., Suzuki, H., Sugimoto, Y. & Sugiyama, Y. ABCG2 Transports Sulfated Conjugates of Steroids and Xenobiotics*. J Biol Chem 278, 22644–22649 (2003).

45. Kurmangaliyev, Y. Z., Yoo, J., LoCascio, S. A. & Zipursky, S. L. Modular transcriptional programs separately define axon and dendrite connectivity. Elife 8, e50822 (2019).

46. Alyagor, I. et al. Combining Developmental and Perturbation-Seq Uncovers Transcriptional Modules Orchestrating Neuronal Remodeling. Dev Cell 47, 38–52.e6 (2018).

47. Uyehara, C. M. & McKay, D. J. Direct and widespread role for the nuclear receptor EcR in mediating the response to ecdysone in Drosophila. Proc National Acad Sci 116, 9893–9902 (2019).

48. Syed, M. H., Mark, B. & Doe, C. Q. Steroid hormone induction of temporal gene expression in Drosophila brain neuroblasts generates neuronal and glial diversity. Elife 6, e26287 (2017).

49. Altmann, C. R. & Brivanlou, A. H. Neural patterning in the vertebrate embryo. Int Rev Cytol 203, 447–482 (2001).

50. Briscoe, J. & Small, S. Morphogen rules: design principles of gradient-mediated embryo patterning. Development 142, 3996–4009 (2015).

51. Gaunt, S. J. Hox cluster genes and collinearities throughout the tree of animal life. Int J Dev Biol 62, 673–683 (2018).

52. Prezioso, G., Giannini, C. & Chiarelli, F. Effect of Thyroid Hormones on Neurons and Neurodevelopment. Horm Res Paediat 90, 73–81 (2018).

53. Miranda, A. & Sousa, N. Maternal hormonal milieu influence on fetal brain development. Brain Behav 8, e00920 (2018).

54. Akin, O. & Zipursky, S. L. Activity regulates brain development in the fly. Curr Opin Genet Dev 65, 8–13 (2020).

55. Ting, C.-Y. et al. Photoreceptor-Derived Activin Promotes Dendritic Termination and Restricts the Receptive Fields of First-Order Interneurons in Drosophila. Neuron 81, 830–846 (2014).

56. Mizumoto, K. & Shen, K. Two Wnts Instruct Topographic Synaptic Innervation in C. elegans. Cell Reports 5, 389–396 (2013).

57. Umemori, H., Linhoff, M. W., Ornitz, D. M. & Sanes, J. R. FGF22 and Its Close Relatives Are Presynaptic Organizing Molecules in the Mammalian Brain. Cell 118, 257–270 (2004).

## Additional References

58. Picelli, S. et al. Full-length RNA-seq from single cells using Smart-seq2. Nat Protoc 9, 171–181 (2014).

59. Buenrostro, J. D., Giresi, P. G., Zaba, L. C., Chang, H. Y. & Greenleaf, W. J. Transposition of native chromatin for fast and sensitive epigenomic profiling of open chromatin, DNA-binding proteins and nucleosome position. Nat Methods 10, 1213–1218 (2013).

60. Ambrosini, G., Groux, R. & Bucher, P. PWMScan: a fast tool for scanning entire genomes with a position-specific weight matrix. Bioinformatics 34, 2483–2484 (2018).

61. Butler, A., Hoffman, P., Smibert, P., Papalexi, E. & Satija, R. Integrating single-cell transcriptomic data across different conditions, technologies, and species. Nat Biotechnol 36, 411–420 (2018).

62. Xu, S. et al. Interactions between the Ig-Superfamily Proteins DIP-α and Dpr6/10 Regulate Assembly of Neural Circuits. Neuron 100, 1369–1384.e6 (2018).

